# Flexible seed size enables ultra-fast and accurate read alignment

**DOI:** 10.1101/2021.06.18.449070

**Authors:** Kristoffer Sahlin

**Affiliations:** Department of Mathematics, Science for Life Laboratory, Stockholm University, 106 91, Stockholm, Sweden

**Keywords:** read mapping, Illumina short-reads, alignment, strobemers, syncmers, seed-and-extend

## Abstract

Read alignment to genomes is a fundamental computational step used in many bioinformatic analyses, and often, it is the computational bottleneck. Therefore, it is desirable to perform the alignment step as fast as possible without compromising accuracy. Most alignment algorithms consider a seed-and-extend approach, where the time-consuming seeding step identifies and decides on candidate mapping locations. Recently, several advances have been made on seeding methods for fast sequence comparison.

We combine two such methods, syncmers and strobemers, in a novel seeding approach for constructing dynamic-sized fuzzy seeds and implement the method in a short-read aligner, strobealign. Firstly, we show that our seeding is fast to construct and effectively reduces repetitiveness in the seeding step using a novel metric E-hits. Secondly, we benchmark strobealign to traditional and recently proposed aligners on simulated and biological data and show that strobealign is several times faster than traditional aligners such as BWA and Bowtie2 at similar and sometimes higher accuracy while being both faster and more accurate than more recently proposed aligners. Our aligner can free up substantial time and computing resources needed for read alignment in many pipelines.

**Availability:** https://github.com/ksahlin/strobealign.

## Introduction

Aligning Illumina sequencing reads to a reference genome is the first step in many analyses pipelines. Due to the fundamental role of short-read alignment in bioinformatics there has been considerable work in this area. BWA (1, 2), Bowtie (3), and Bowtie2 (4), which use the Burrows-Wheeler transform (5), have received widespread use through their favourable memory and runtime trade-off. These aligners have for several years been the dominating genomic short-read aligners. Several alternative approaches have been proposed, such as random permutations aligner (6), rNA (7), SNAP (8), and subread (9) that has alternative strengths showing improved accuracy or decreased runtime in specific use cases. A comprehensive listing of alignment techniques is found in (10). There are also several major contributions in data structures and algorithms in splice alignment of RNA-seq data (11, 12) or pseudo-alignment methods (13, 14) that are incredibly fast by not explicitly performing an exact alignment.

A pioneering approach was the use of minimizers (15, 16) as seeds in alignment and overlap detection algorithms (17, 18). Minimap2 was initially described for long-read alignment but has shown similar alignment accuracy of short reads as popularly used BWA and Bowtie2. Minimap2 uses minimizers as seeds and then employs collinear chaining to produce candidate locations for the alignment step. Along the same trajectory, Mashmap (19) and Winnowmap (20) were designed for long-reads and made algorithmic contributions to minimizer-based alignment by considering and adjusting the densities and probabilities around the sampling of minimizers.

A computational hurdle in sequence alignment is the length of the seeds, which inform the aligner of candidate mapping locations. Alignment algorithms often need to use a shorter seed length than what gives unique matches in a genome to have good alignment accuracy. Therefore, seeds may produce many candidate regions that need to be filtered based on some score before the alignment stage. Therefore, alignment methods are usually described to employ a seed-filter-extend approach, where the seeding and filtering are at the heart of an aligner’s performance. In (21), the authors propose an alternative approach to the filtering step by computing a much cheaper Hamming distance of an embedded representation of the read. Interesting candidate sites should have low embedded Hamming distance and are sent for alignment. Their aligner, Accel-Align, outperforms other aligners in terms of speed.

While filtering is an important step for candidate selection, reducing the false matches caused by repetitive seeds is preferable. To do so, one has to increase seed size. In this work, we describe a seeding approach for creating seeds with variable sizes that are fuzzy, i.e., they can match over mutations. Our approach allows us to use much longer seed lengths than other approaches without loss in accuracy. Furthermore, longer seed lengths reduce the number of candidate sites to evaluate, allowing much faster mapping and alignment. Our contribution is based on two recent advancements in the area of sequence comparison; syncmers (22) and strobemers (23). Both syncmers and strobemers have been demonstrated to improve sequence similarity detection (23, 24). Syncmers were proposed as an alternative to the minimizer subsampling technique. In contrast, strobemers were proposed to produce gapped seeds as an alternative to *k*-mers and spaced *k*-mers.

Here we show that syncmers and strobemers can be combined in what becomes a high-speed indexing method, roughly corresponding to the speed of computing minimizers. Our technique is based on first subsampling *k*-mers from the reference sequences by computing canonical open syncmers (22), then producing strobemers (23) formed from linking together syncmers occurring close-by on the reference using the randstrobe method. A consequence is that instead of using a single seed (e.g., *k*=21 as default in minimap2 for short-read mapping), we show that we can link together two syncmers as a strobemer seed and achieve similar accuracy to using individual minimizers.

Our first contribution in this work is the novel seeding approach by computing strobemers over syncmers. We demonstrate that this seeding is fast to compute (less than 5 minutes to index hg38) and is therefore competitive to, *e.g*., minimizers in high-performance sequence mapping scenarios. Our seeding method constitutes a novel algorithm class according to the comprehensive aligner classification table in (10), with our method being hashing of variable-length fuzzy seeds. We further evaluate the seed-repetitiveness of our seeding compared to some of the current approaches in read mappers (*k*-mers, minimizers, syncmers) using a novel metric E-hits, and show that our seeds are effective at reducing seed-repetitiveness. We believe that E-hits will be a useful metric for further development and comparisons of seeding approaches.

Our second contribution is implementing our seeding technique in a short-read alignment tool, strobealign. We use simulated and biological data to show that strobealign is several times faster than traditional aligners such as BWA and Bowtie2 while being faster and more accurate than more recently proposed aligners. Interestingly, we observe that sub-sampling methods such as strobealign and minimap2 can also be more accurate than BWA-MEM on high diversity datasets. Strobealign reaches peak performance on paired-end reads of lengths 150-300 nucleotides (nt), which is well suited for advances in short-read sequencing length and throughput. An example of such an advance is Illumina’s Chemistry X reads that are claimed to be two times longer (25). Strobealign’s speed can remove the alignment bottleneck and free up substantial computing resources in many pipelines.

## Results

### Method overview

We present a seeding method for sequence similarity search that is based on a combination of two previously published techniques syncmers (22) and strobemers (23). The main idea of the seeding approach is to create fuzzy seeds by first computing open syncmers from the reference sequences, then linking the syncmers together using the randstrobe method (23) with two syncmers (Fig. 1A). Our fuzzy seeds enable us to use larger seed lengths that are more likely to be unique (concept illustrated in Fig. 1B) while still allowing mutations or read errors between the syncmers. Figure 1C illustrates the seeds extracted from a DNA sequence.

**Fig. 1.**
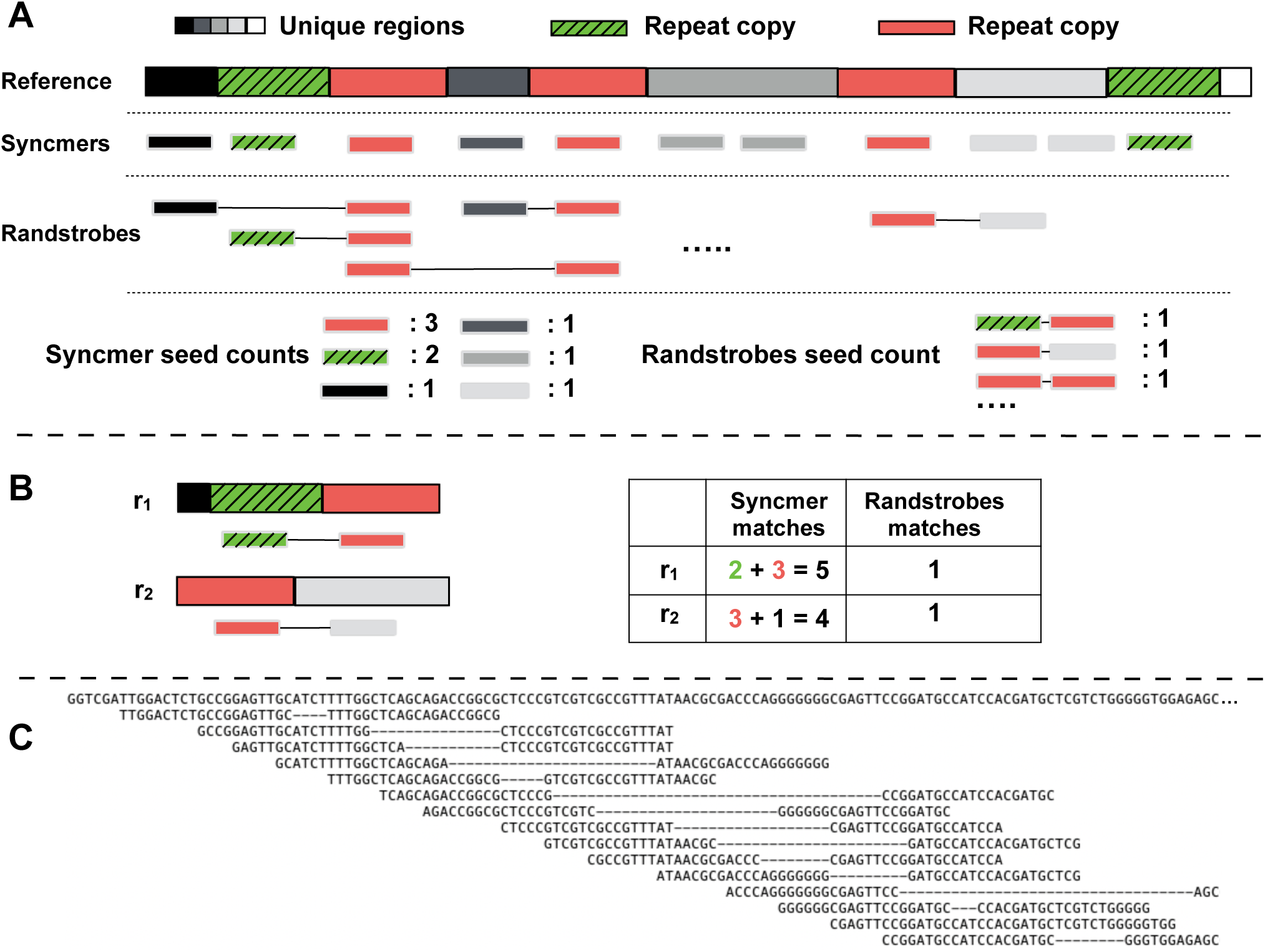
Overview of seeding construction. (A) Open syncmers are constructed from the sequence, and randstrobes are constructed by linking together syncmers. The second strobe is sampled with a minimum and maximum spread between [*w*_*min*_, *w*_*max*_] strobes downstream from the first strobe. While syncmers may occur several times due to repeats on the genome (red and green boxes), randstrobes are less repetitive. (B) Two reads, *r*_1_ and *r*_2_, are mapped to the reference. Finding matches using only syncmers creates several candidate mapping locations, while the randstrobes are unique in this scenario. In the illustration, the syncmers are spread out for visual purposes. Panel C shows a real sampling of randstrobes on a DNA sequence and their spread using (2, 20, 4, 11) with sampling skew.

Compared to the study introducing strobemers (23), we present two novel ideas. Firstly, (23) described strobemers as linking together strobes in ‘sequence-space’, *i.e*., over the set of all *k*-mers. Since syncmers represent a subset of *k*-mers from the original sequence, computing randstrobes over this subset of strings is very fast while still having a similar linking range over the original sequence. Secondly, we also introduce a skewed linking approach of the second strobe that links nearby strobes more often (Suppl. Fig. S1) which is more effective for shorter read lengths than the original approach (23) (details in the method section). We implement our seeding approach as the fundamental technique in a new short-read aligner strobealign.

### Overview of read-alignment evaluation

#### Tools

We evaluated strobealign (v0.7.1) to six state-of-the-art and recently published short-read aligners, BWA-MEM (v0.7.17), BWA-MEM2 (v2.0pre2), Bowtie2 (v2.3.5), minimap2 (v2.22), Accel-Align (v1.1), SNAP (26) (v2.0.0) on simulated and biological Illumina sequencing data sets. We also attempted to evaluate URMAP (27) (v1.0.1480_i86linux64) and PuffAligner (28) but were unable to run them for our experiment designs (see Suppl. Note C). We ran all the tools using 16 threads as multithreading is the standard use case. We also investigated how the tools scaled with the number of threads by running them with 4, 8, and 16 threads in one analysis. Since BWA-MEM and BWA-MEM2 have identical accuracy, we will only refer to BWA-MEM when discussing accuracy (to imply BWA-MEM and BWA-MEM2) but evaluate them separately in terms of alignment time and memory.

#### Simulated data

We simulated single-end and paired-end reads with read lengths of 50, 75, 100, 150, 200, 250, 300, and 500 nucleotides. We chose these lengths as 50 and 75 are used in applications such as for chromatin profiling or RNA-seq quantification analyses, while 100 to 250 are within the range of standard Illumina protocol read lengths. To investigate performance on future Illumina chemistry X reads, which are claimed to be two times longer than current protocols, we simulated read lengths of 300 and 500 to emulate read lengths of two times the 150 and 250 protocols, respectively. While we include read lengths of 50 and 75nt, we emphasize that strobealign is designed for Illumina read lengths 100nt and above typically used, *e.g*., for genomic alignment and downstream SNP and indel calling and similar scenarios. In our first simulated experiment, we simulated ten million single-end and paired-end reads from human genomes at four different divergence levels from hg38 denoted SIM1, SIM2, SIM3, and SIM4, where SIM4 has the highest divergence from hg38 (details on simulations in Suppl. Note A).

In our second simulated experiment, we simulated ten million paired-end reads from the genomes of the fruit fly (180Mb), maize (2.4Gb), human cell line CHM13 (29) (3.2Gb), and rye (30) (7.3GB), denoted drosophila, maize, CHM13, and rye, respectively, where reads were simulated at the same diversity level as for the SIM3 dataset. For details on this experiment, see Supplementary Note A. These genomes are of variable sizes, with the latter three more repetitive than hg38. In our third simulated experiment, we evaluated the aligners by simulating ten million paired-end reads with high SNP and indel rate from a simulated repetitive genome (denoted REPEATS). The REPEATS genome consisting of five hundred 100kbp copies at roughly 90% similarity (details on simulations in Suppl. Note A).

In our fourth and final simulated experiment, we simulated two larger paired-end read datasets with read lengths 150nt and 250nt denoted SIM150 and SIM250, respectively. The SIM150 and SIM250 datasets contain 300M and 180M reads, respectively, each constituting roughly 30x coverage of hg38. The reads were simulated from a genome with a SIM3 variation rate to hg38. We used these two datasets to evaluate downstream SNP and indel calling from the alignments, and they were chosen to match the lengths of our biological data (described in the next section).

#### Biological data

We evaluated downstream SNP and indel calling results on two biological paired-end Illumina read datasets of lengths 150nt and 250nt from HG004 in the Genome-In-a-Bottle consortium (31). We denote these datasets BIO150 and BIO250. The BIO150 and BIO250 datasets have a rough coverage of 32 and 26, respectively. Finally, we also used a subset of 4M reads from BIO150 and BIO250 to evaluate the mapping concordance between the aligners. Details of the datasets are found in Suppl. Note D and details of the SNP and indel calling pipeline in Suppl. Note E.

#### Evaluation metrics

We will use the following terminology. With a read mapping, we mean to find the location of a read on the reference. With read alignment, we mean that the read is not only assigned a location but has also been pairwise aligned to the reference at the mapping site. We evaluated the mapping accuracy of aligners by looking at the overlap of read alignments with the correct genomic location. For nucleotide level accuracy (such as aligning in and around indels), we evaluate the downstream SNP and indel calling results that the alignments produce. We also evaluate the runtime and memory usage of each aligner. For details on how the evaluation metrics are computed, see Suppl. Note B. For the aligners that also support a mapping mode (Accel-Align, minimap2, and strobealign), we evaluated both the mapping and alignment modes (details on running aligners in Suppl. Note C).

### Seeding results

An important factor for fast and accurate mapping is that the seeds are relatively unique on the reference genome. We compared two metrics (E-hits and fraction hard masked seeds) related to seed uniqueness. E-hits indicates how many spurious hits to the reference are found on average, given that reads are drawn uniformly at random from the reference genome. A formal definition of E-hits is given in subsection the E-hits metric in Methods. The fraction of hard-masked seeds is the fraction of seeds with occurrence over 1000 times in the reference and are excluded from seed finding (see section Implementation details).

We compared E-hits and fraction hard masked seeds for different lengths of *k*-mers, minimizers with density 1/5 (w=9), open syncmers with density 1/5 (s = k-4, t=20), and strobealign seeds for the various read lengths (labeled sa*X, X* ∈ { 50, 75, 100, 150, 200, 250, 300, 500 } (all with 1/5 in density except sa500 that has a density of 1/7). Since strobealign seeds are flexible in seed size, we use the median seed size in our analysis. We used jellyfish (32) to obtain the *k*-mer counts and a custom script (provided in Data Availability) to obtain the minimizers and syncmers. All seeds are canonically represented (smallest seed hash value out of forward and reverse complement is stored) as is standard in read alignment (details in section Modifications to strobemers).

We first observe that subsampling minimizers and syncmers will create a more repetitive index than using all k-mers for seeds of length 20-30, but evens out as seeds become longer (Fig. 2A-B). This is relevant to read mapping since most aligners use seeds of around 20-30nt. Secondly, we see that syncmers produce more repetitive index than minimizers, as has been proven in (24).

**Fig. 2.**
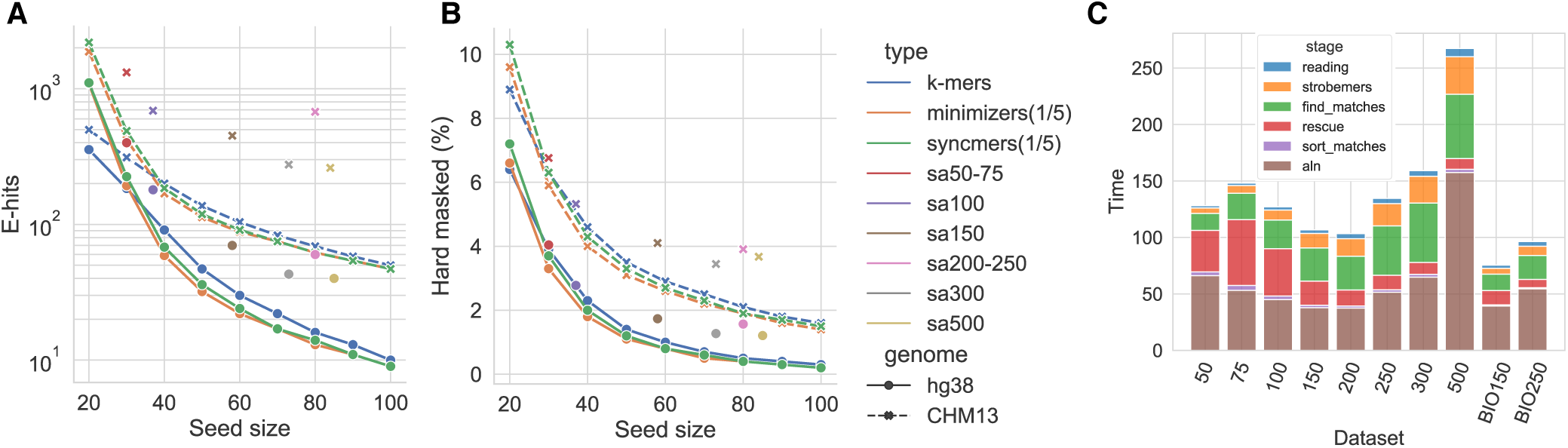
Seed uniqueness and time allocation in strobealign. Panel **A** shows the expected number of hits from a seed randomly drawn from reference (E-hits) for some popular seeding approaches (*k*-mers, minimizers, syncmers) in comparison to strobealign’s seeds. Minimizers and syncmers are both sampled at a sub-sample fraction of 1/5, and minimizers use a random hash function. For strobealign’s seeds which have variable lengths, the median seed length is plotted. Strobealign’s seeds for read lengths 100-500nt (typically two linked syncmers of length 20) reduce the repetitiveness an order of magnitude on hg38 compared to using a single syncmer or minimizer of length 20. Panel **B** shows the fraction of seeds that would be hard masked in strobealign (occurring over 1000 times). On hg38, strobealign’s seeds for read lengths 100-500nt hard masks 2.6-6 times fewer seeds over syncmers of length 20. Panel **C** shows the real time spent at various steps in strobealign using 16 threads for the SIM3 datasets of 10 million paired-end reads of different read lengths and the subsampled BIO150 and BIO250 datasets of 4 million paired-end reads of length 250. Reading refer to reading the fastq files. The label ‘strobemers’ refers to the time to generate strobemer seeds from the reads, ‘find_matches’ refers to retrieving and creating merged matches from all strobemer seeds below the repetitive abundance threshold, ‘rescue’ refers to finding merge matches in the rescue mode, ‘sort_matches’ sorts the matches with respect to the candidate map score, and ‘aln’ refers to the base level alignment, in which the large majority of runtime constitutes of calling ssw and a small fraction is computing the hamming distance. Writing the output to SAM was not logged in the experiments but typically takes less time than reading input.

When studying how strobealign’s seeds fare to exact seed approaches, we observe that strobealign’s seeds, which use two syncmers of length about 20 (see Implementation details), reduce the repetitiveness over using only one syncmer of length 20. For example, when mapping reads of length 150nt (sa150), which pairs two syncmers of length 20nt, strobealign’s seeds achieve over 15 times lower E-hits than using only one syncmer of 20. The sa150 seeds are also more than three times less frequent to be hard masked on hg38 (Fig. 2B). The sa150 seeds are comparable to syncmers of size 40 in E-hits and between 40-50 in the fraction of hardmasked seeds.

When comparing sa150 seeds to *k*-mers, the sa150 seeds are as unique as using *k*-mers of about 45 (Fig. 2A-B). Such long solid *k*-mers are not suited for short-read alignment, and previous studies have typically decided on sizes between 20 and 32 (2, 18, 21, 27). For the 300 and 500nt reads, strobealign’s seeds have roughly the same statistics as *k*-mers of length 55. On CHM13, the seeding results show a similar trend, although not as strong. Here, sa150 only archives the same E-hits score as using k-mers slightly larger than 20. However, sa150 has much lower hard masking, comparable to k-mers of about size 45.

In summary, our study of E-hits and the fraction of hard masked seeds highlights two points. Firstly, our seeds can achieve the same uniqueness (E-hits and fraction hard masked) as k-mers with lengths traditionally unsuitable for short-read alignment. Secondly, when constructing strobealign seeds, the linking process (strobemers) is responsible for the major reduction in repetitive seeds, as can be seen by comparing strobealign’s seed uniqueness to only using a single minimizer or syncmer of length of about 20 (similarly to what is done in minimap2).

### Indexing results

We measured the time and memory to produce our dynamic-sized seeds on five genomes; drosophila, maize, human (hg38 and CHM13), and rye (Table 1). Our seeding is relatively fast. For example, on hg38, while the total indexing time is 259 and 167 for strobealign and minimap2, respectively, it takes only 133 seconds to construct strobealign’s seeds, compared to 100 seconds to produce minimizers in minimap2. The remaining indexing time is spent on sorting the seeds (standard library sort in C++) and creating a hash table. These are steps that can be further optimized in strobealign by changing algorithms, *e.g*., as using radix sort as in minimap2. Our indexing is also faster than most of the other aligners (Table 1). Furthermore, the seeding is not a bottleneck in the alignment step, taking up only a small fraction of the total alignment runtime across datasets (yellow segment in Fig. 2C). As for the peak memory, strobealign has a peak indexing memory footprint of about 31Gb in hg38 and 50Gb on rye, placing it in fourth place behind BWA-MEM, Bowtie2, and minimap2.

**Table 1.** Indexing time and peak memory of indexing for aligners using one thread. Strobealign’s indexing time and memory depends on the syncmer density used. ^1^ When using syncmer density 1/5 (50-300nt datasets). ^2^When using syncmer density 1/7 (500nt datasets). ^3^ SNAP could not index the rye genome (Suppl Note. C). ^4^AccelAlign does not have a singe thread mode for indexing. Multithreaded results are displayed on a node with 20 cores. We observed it used 300-700% CPU during indexing.

|  | Drosophila |  | Maize |  | hg38 |  | CHM13 |  | Rye |  |
| --- | --- | --- | --- | --- | --- | --- | --- | --- | --- | --- |
|  | Time (s) | Mem (Gb) | Time (s) | Mem (Gb) | Time (s) | Mem (Gb) | Time (s) | Mem (Gb) | Time (s) | Mem (Gb) |
| minimap2 | **9** | 1.3 | **125** | 9.8 | **167** | 13.0 | **191** | 13.2 | **421** | 24.8 |
| Strobealign<sup>1</sup> | 12 | 3.4 | 176 | 15.3 | 259 | 31 | 268 | 31.5 | 598 | 50 |
| Strobealign<sup>2</sup> | **9** | 2.3 | 144 | 15.0 | 210 | 22.2 | 222 | 22.6 | 545 | 44.4 |
| SNAP | 63 | 1.9 | 1210 | 38.6 | 1,744 | 45.9 | 1826 | 48.0 | NA<sup>3</sup> | NA<sup>3</sup> |
| BWA-MEM | 120 | **0.2** | 2,397 | **3.2** | 3,684 | **4.5** | 3,629 | **4.6** | 10,245 | **10.7** |
| BWA-MEM2 | 99 | 4.2 | 1,365 | 63.9 | 3,146 | 90.8 | 3,000 | 91.3 | 7,092 | 212.2 |
| Bowtie2 | 228 | 0.3 | 5629 | 4.1 | 6,008 | 5.5 | 7,002 | 5.8 | 24,756 | 20.0 |
| AccelAlign<sup>4</sup> | 10 | 3.4 | 101 | 28 | 134 | 38.8 | 132 | 40 | 320 | 92.5 |

**Table 2.**
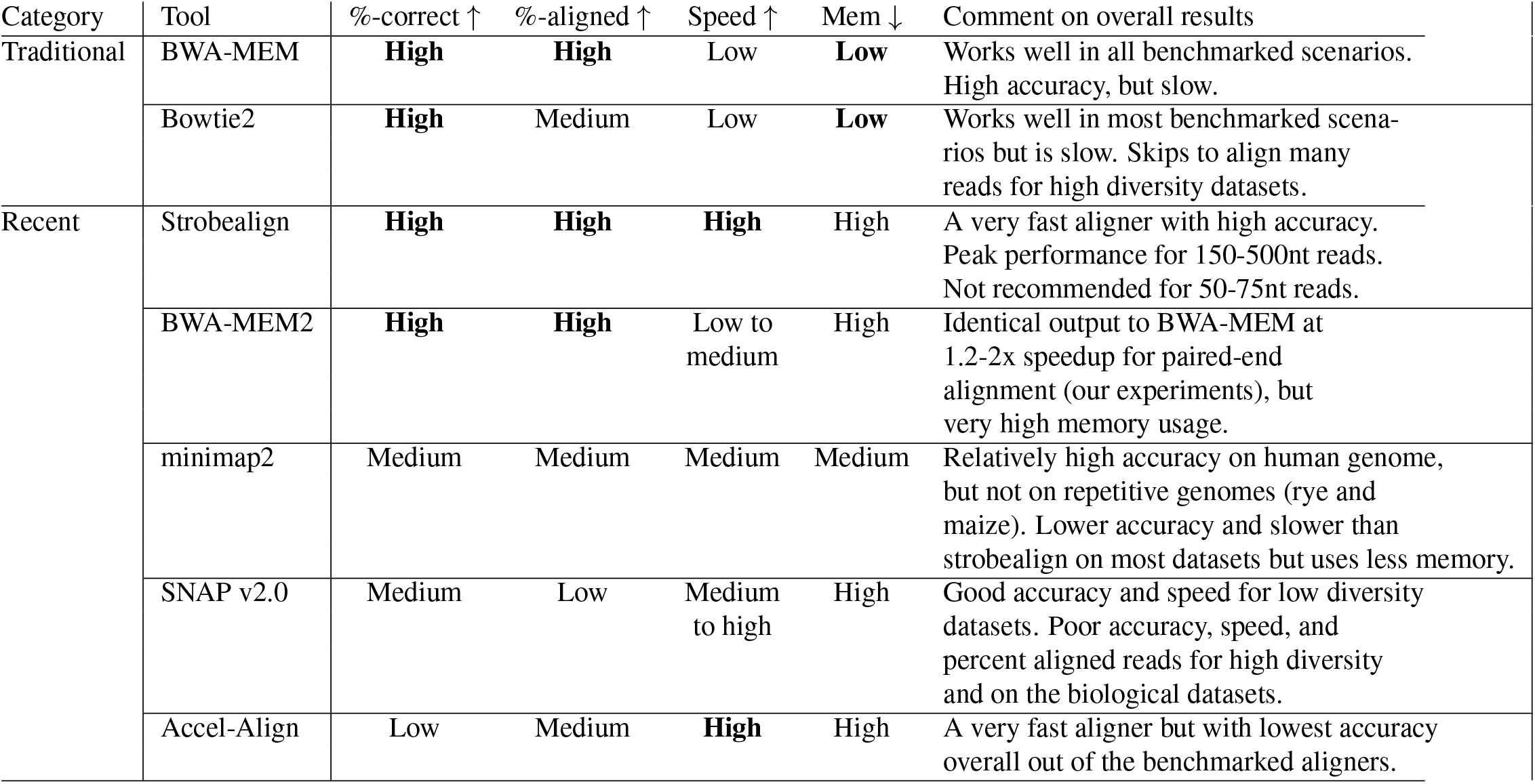
Overview of aligner characteristics based on results from our experiments. Brief comments on the characteristics of the aligners are included.

### Results on hg38 simulated data

For the paired-end read experiments, we present the accuracy, runtime, and percent aligned reads for the SIM3 dataset (Fig. 3), which has a SNP and small indel rate roughly observed in a human population. The remaining three datasets with both lower (SIM1 and SIM2) and higher (SIM4) variation rates are found in Suppl. Fig. S2-4. When looking at alignment accuracy for the shortest read lengths of 50-75nt that are useful in experiments such as RNA-seq gene expression profiling or chromatin profiling, the traditional aligners BWA-MEM, BWA-MEM2 and Bowtie2 have the highest accuracy. For these types of analyses and read lengths, BWA-MEM, Bowtie2, or specialized aligners such as Chromap (33) for chromatin data or pseudoalignment methods such as Kallisto (13) or Salmon (14) for RNA-seq reads should be used. Since strobealign is currently not designed for this type of data, we, from now on, focus our evaluation on the common genomic analysis read lengths of 100-250nt, and future chemistry X read lengths of 300 and 500.

**Fig. 3.**
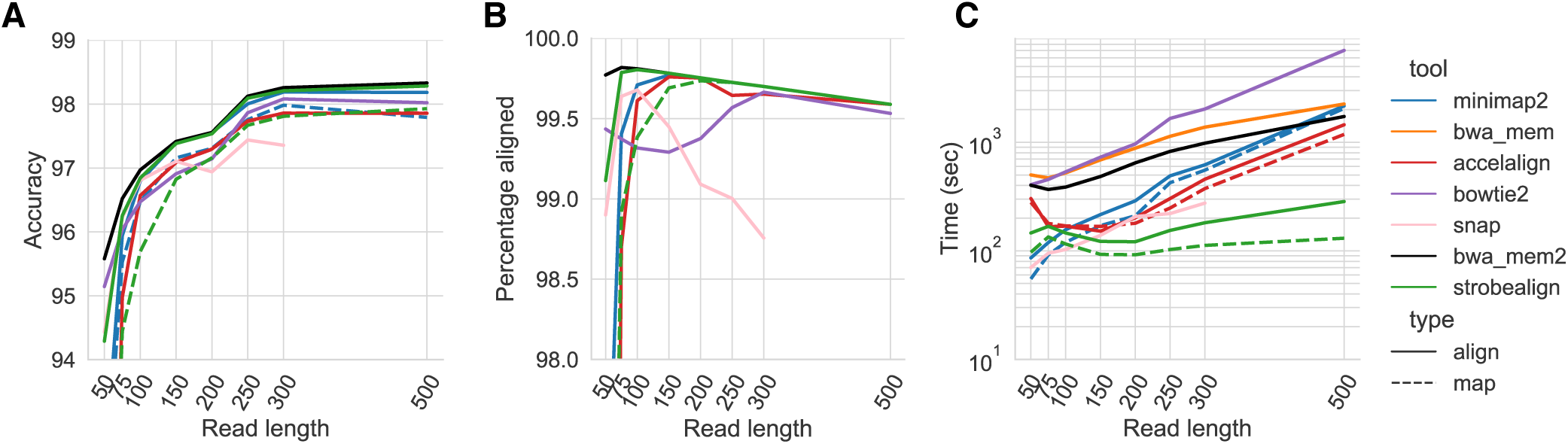
Accuracy (Panel A), percent aligned reads (Panel B), and runtime (Panel C) of aligning paired-end reads to the SIM3 dataset.

For SIM1-SIM3 with read lengths 150nt and above, strobealign has at most 0.1% lower accuracy than BWA-MEM (read lengths 300 and 500 on SIM2; Suppl. Fig. S2), but the accuracy gap is only 0.05% to non-existent on SIM3 (Fig. 3A). For SIM4, strobealign has higher accuracy than BWA-MEM with about 0.1%. Overall, strobealign and BWA-MEM typically have the highest and second-highest accuracy on most of the 150-500nt datasets, although other aligners achieve good accuracy on individual datasets. Specifically, minimap2 has the highest accuracy on SIM4 for read length 150 and 200 (about 0.05% higher than strobealign), and SNAP and Bowtie2 has high accuracy on the low diversity datasets SIM1 and SIM2. While SNAP’s accuracy is high for the low diversity datasets, it quickly becomes non-competitive for higher diversity (Fig. 3A, Suppl. Fig. S2). For read lengths of 100nt, BWA-MEM has 0.15% higher accuracy than strobealign across the SIM1-SIM4 datasets. Strobealign and BWA-MEM are typically also able to align the most reads (Fig. 3B, Suppl. Fig. S4).

As for alignment time for read lengths of 100-250nt, strobealign is about 7 times faster than BWA-MEM and 4.5-6 times faster than BWA-MEM2 (Fig. 3C, Suppl. Fig. S3). SNAP also shows competitive runtime on the low diversity datasets SIM1 and SIM2 (Suppl. Fig. S3). For the simulated chemistry X read lengths of 300nt and 500nt, strobealign is 6.4-9 times faster than BWA-MEM and 4.5-7 times faster than BWA-MEM2 (Fig. 3C, Suppl. Fig. S3).

In addition to being several times faster than BWA-MEM2, strobealign also uses 1.5-2.1x lower peak memory than BWA-MEM2 (Suppl. Fig. S5). BWT-based aligners such as BWA-MEM and Bowtie2 use much lower peak memory than all other aligners (Suppl. Fig. S5), where Bowtie2 has the lowest peak memory across all experiments in this analysis.

We also ran the alignment tools with 4, 8, and 16 threads on the different datasets from the SIM3 dataset. Alignment time nearly halves for the tools when doubling the number of threads suggesting that the tools utilized the resources well (Suppl. Fig. S7). Relative alignment times between the tools for 4 and 8 threads stay similar to our benchmarks using 16 threads.

For our single-end data experiments (Suppl. Fig. S8-11), we largely observe the same results reported for the paired-end evaluation. Strobealign is the fastest tool for all read lengths of 150nt and above. Although strobealign has slightly worse performance on high diversity datasets in single-end mode (panels SIM3 and SIM4 in Suppl. Fig. S8), it has substantially higher accuracy and percentage of aligned reads to the tools with similar speed (Accel-Align and SNAP). Minimap2 performs well for the single-end reads of the highest diversity (SIM4). The analysis is reported in detail in Suppl. Note F. Finally, as expected, the mapping modes of minimap2 and strobealign have lower accuracy than their respective alignment modes. However, strobealign’s mapping mode has a more substantial accuracy reduction to its alignment mode than minimap2. This indicates that there might be room to develop a better strobemer-based scoring function that more frequently assigns the true mapping location with the highest score. However, the primary purpose of short read alignment is that alignment and mapping mode is not implemented in many aligners.

In summary, for the hg38 paired-end read analysis, strobealign has the best tradeoff between accuracy, runtime, and percent aligned reads to any of the other benchmarked aligners on most of the datasets. Strobealign and BWA-MEM have the highest accuracies across diversity levels for reads of lengths 150nt and above and are usually within a difference of 0.1% to each other. On the high diversity dataset SIM4 for read lengths of 150nt and longer, there is no tradeoff between accuracy and runtime between the two tools, as strobealign is several times faster and more accurate than BWA-MEM and BWA-MEM2, as well as uses lower memory than BWA-MEM2. A notable aligner is SNAP, which has high accuracy and is very fast on SIM1. However, its performance across accuracy, speed, and percentage of aligned reads deteriorates substantially with increased diversity (Fig. 3, Suppl. Fig. S2).

### Results on other genomes

Our benchmarks on the four additional genomes drosophila, maize, CHM13, and rye broadly show similar results to our experiments on hg38. That is, on most datasets with read lengths 150nt or longer, strobealign and BWA-MEM have substantially higher accuracy than the other aligners (Suppl. Fig. S12). For example, strobealign is slightly more accurate than BWA-MEM (about 0.05%) on drosophila and slightly less accurate (at most 0.11%) than BWA-MEM on the new human genome CHM13 (Suppl. Fig. S12C). In addition, Strobealign is consistently 7-9 times faster than BWA-MEM on the maize, CHM13, and rye genomes (Suppl. Fig. S13) and 4-5 times faster than BWA-MEM2 and uses 2-3 times less peak memory than BWA-MEM2 (Suppl. Fig. S14).

On maize and rye, strobealign does not reach comparable accuracy with BWA-MEM for read lengths of 150 and 200 (Suppl. Fig. S12). All aligners, including strobealign, have been run using default parameters which may not be optimal for particular read lengths or genomes. To study if we could reduce this gap, we specified the parameter -M 40 to strobealign to consider more alignment locations (default is 20). With this setting, we observed that the gap in accuracy nearly disappeared on maize and was reduced on rye (Suppl. Fig. S16), while alignment time was still about 5-7.5 times faster than BWA-MEM and about 3-3.5 times faster than BWA-MEM2 (Suppl. Fig. S17). With -M 40, strobealign’s accuracy also remains close to identical to BWA-MEM on drosophila and CHM13 while being, *e.g*., around 7 times faster than BWA-MEM and 4-4.5 times faster than BWA-MEM2 on CHM13 (Suppl. Fig. S16-17).

As for general memory usage (Suppl. Fig. S14), strobealign’s indexing scales with the number of unique seeds. For example, Strobealign uses only 1.5 times more memory for a genome that is 2.3 times as large (rye). This scaling is not observed in BWA-MEM2. As for the number of aligned reads, BWA-MEM and strobealign align the most reads in general (Suppl. Fig. S15).

Finally, we observe that many tools face complications when aligning to rye. Minimap2’s percentage of aligned reads and accuracy drop to less than 50% accuracy on most instances and does not show in the figures. Also, SNAP and Accel-Align could not run on this dataset (Suppl. Note C).

### Results on the REPEATS dataset

We ran the aligners in paired-end mode on the REPEATS dataset, which constitutes a particularly challenging repetitive dataset with high diversity (described in Suppl. Note A). On this dataset, strobealign and minimap2 have the highest accuracy (Fig. 4A). However, minimap2’s accuracy comes at the cost of runtime on this dataset, where it is as slow as BWA-MEM or slower. Strobealign has the highest accuracy, most aligned reads, and the fastest runtime for all the read lengths of 150nt and longer ((Fig. 4A-C), being 5-10 times faster than minimap2, 4-6 times faster than BWA-MEM, and about 3.5 times faster than BWA-MEM2. Accel-Align also has a relatively competitive accuracy-runtime tradeoff on the 150nt and 200nt read lengths in this experiment.

**Fig. 4.**
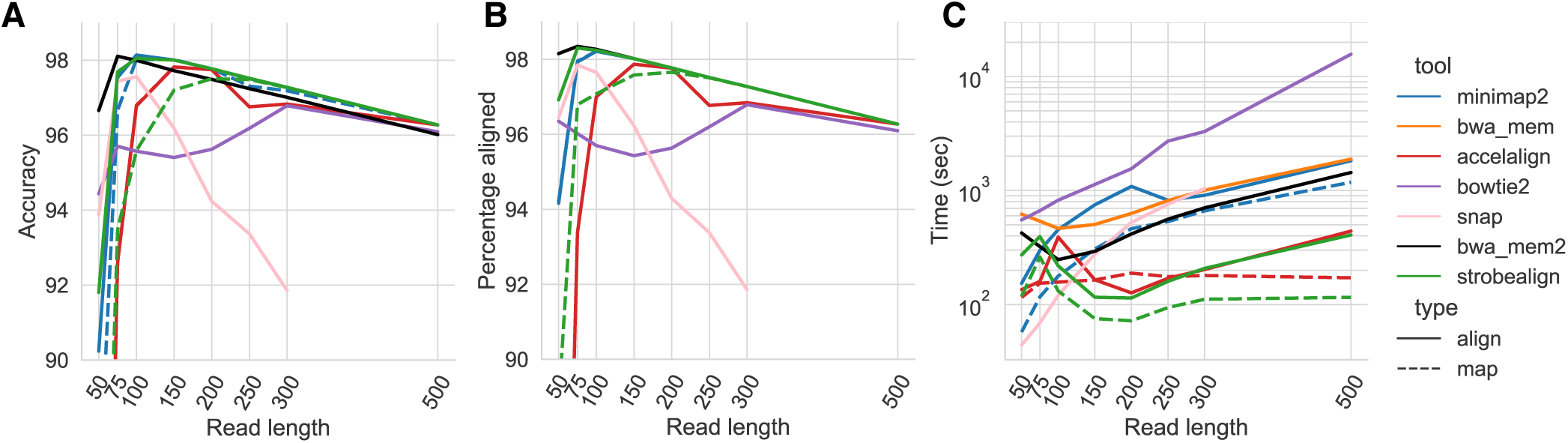
Accuracy (Panel A), percent aligned reads (Panel B), and runtime (Panel C) of aligning paired-end reads to the REPEATS dataset.

### Mapping concordance on biological data

As we do not have the ground truth genomic location of the biological reads, we used a subset of the BIO150 and BIO250 datasets (details in Suppl. Note D) to investigate read mapping concordance. We used the slower but tried-and-tested tools BWA-MEM (BWA-MEM2) and Bowtie2 as gold standard as we observed that they aligned the most reads (Fig. 5A). We then scored the rest of the aligners based on mapping concordance with the two tried-and-tested aligners. Specifically, we measured the concordance in alignment coordinates between three tools; BWA-MEM, Bowtie2, and each of the remaining aligners (Fig. 5B). We assume in this analysis that a high concordance with BWA-MEM and Bowtie2 is a proxy of high accuracy. We also measured runtime and the percentage of aligned reads.

**Fig. 5.**
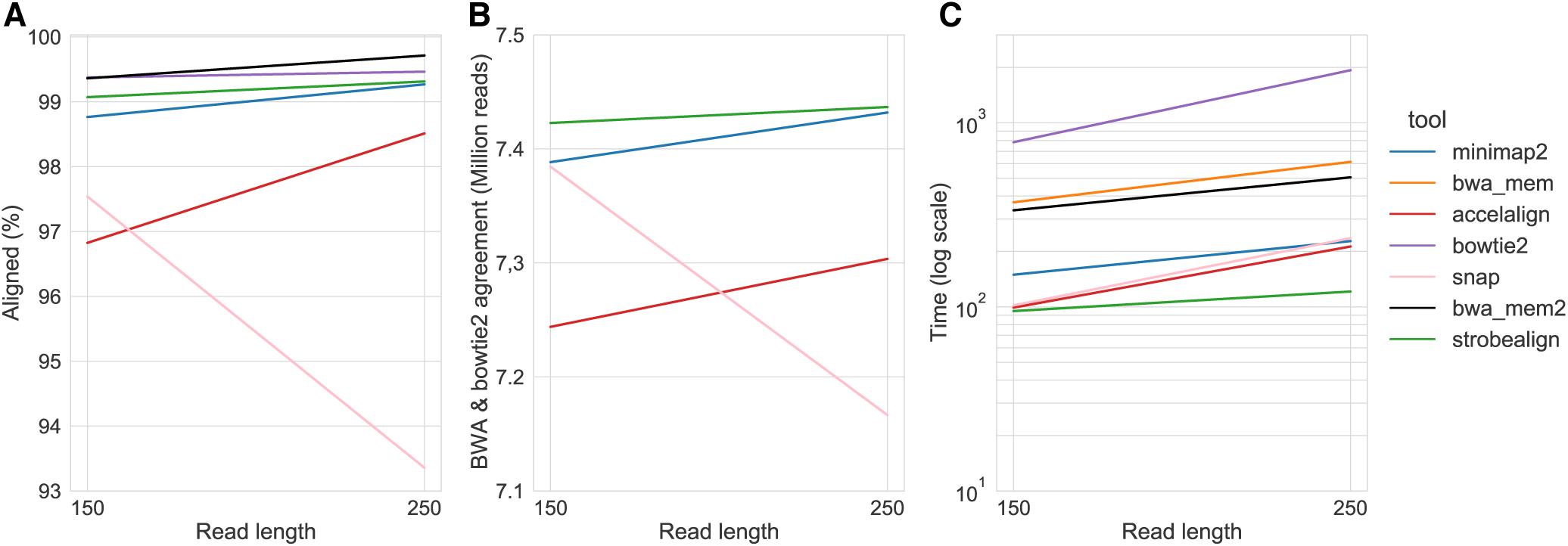
Alignment concordance results for a subset of of 4 million paired-end reads of the BIO150 and BIO250 datasets. Panel A shows the percentage of aligned reads. Panel B shows for strobealign, minimap2, Accel-Align, SNAP, the number of reads that were aligned to the same location (overlapping alignment coordinates) out of the reads that Bowtie2 and BWA-MEM/BWA-MEM2 aligned to the same location (*i.e*., three-way concordance). Panel C shows the runtime.

We observe that strobealign is the fastest tool, has the most aligned reads, and is most consistent with BWA-MEM and Bowtie2 (Fig. 5) for both datasets. The second best performing aligner in this analysis is minimap2. We further looked at the detailed concordance Venn diagrams between BWA-MEM, Bowtie2, strobealign, and minimap2 (Suppl Fig. F12). Minimap2 shares a substantial fraction of alignments with BWA-MEM and relatively few with Bowtie2, while strobealign has a more evenly distributed concordance diagram between Bowtie2 and BWA-MEM. Some of the overlaps we see uniquely shared by two aligners may be on the implementation level by choosing the same random mapping location in case of ties, as aligners have different methodologies to select locations in ambiguous scenarios.

### SNP and indel calling analysis

A common application downstream of read alignment is SNP and indel prediction. While a high mapping accuracy (correct read location) is desired, an aligner also needs accurate base-level alignments when calling SNPs and, in particular, indels. While such an analysis supplements a mapping accuracy analysis, a caveat is that variant callers use MAPQ scores for SNV and indel prediction (34), therefore some callers may be developed or tuned based on popular aligners’ MAPQ scores. Specifically, it has been shown that *bcftools call* was the SNP caller that produced the best result with BWA-MEM alignments out of seven variant calling tools (35). Nevertheless, we used *bcftools call* to benchmark recall, precision, and F-score of SNP and indel calling from the aligners’ output on SIM150, SIM250, BIO150, and BIO250 datasets. Details on the data and analysis pipeline are found in Suppl. Note D and E, respectively.

As for SNP calling, strobealign has 1% and 1.9% lower recall than BWA-MEM on the BIO150 and BIO250 datasets (Fig. 6A; left panel) but has the highest precision out of all aligners, with a 5.4% and 5.3% higher precision than BWA-MEM (Fig. 6A; center panel). When combined, strobealign has the highest SNP calling F-score on both the biological datasets among all aligners with 2.7% and 2.5% higher F-score than BWA-MEM (Fig. 6B; right panel). On the simulated datasets, the recall and precision are similar for all aligners except Accel-Align. Strobealign has the second-highest F-scores after BWA-MEM with an average of 0.45% lower recall at the same precision, resulting in a 0.25% lower F-score than BWA-MEM.

**Fig. 6.**
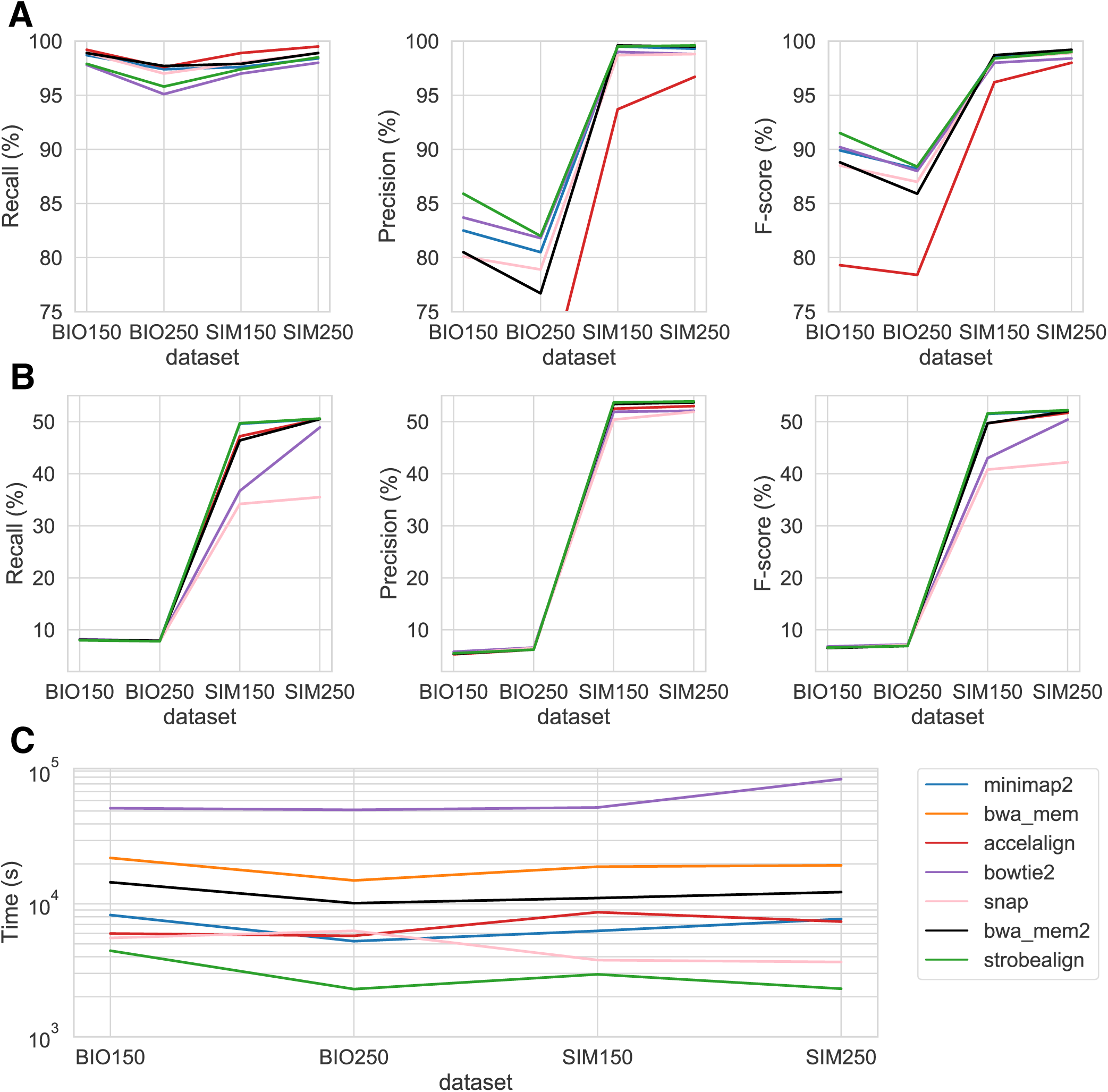
SNP and indel calling with bcftools on SIM150, SIM250, BIO150, and BIO250. Panel A and B shows recall, precision and F-score for SNP and indel calling, respectively. Panel C shows alignment runtime. In panel A, the y-axis was cut at 75% for visibility. Accel-Align has 66.0% and 65.5% for BIO150 and BIO250, respectively.

As for indel calling, which for the biological datasets was computed against all gold-standard variants found by the GIAB consortium, all the aligners have a low (and similar) recall and precision. BWA-MEM has a 0.1% higher recall but a 0.1% lower precision over strobealign on BIO150 and BIO150. However, for the indel calling on simulated data, strobealign has both the highest recall and precision across aligners, with a substantial increase in recall on the SIM150 dataset (3.3%, Fig. 6B), giving the highest F-scores on both the datasets.

As for runtime, strobealign is the fastest aligner across the four datasets, with a 5-8.5 times speedup over BWA-MEM and 3.3-5.3 times speedup over BWA-MEM2.

## Discussion

We have presented a novel approach to compute seeds used for sequence mapping. We showed that our seeding method is fast to construct (Table 1) and that our seeds can achieve the same uniqueness as k-mers with lengths traditionally unsuitable for short-read alignment (Fig. 2A-B).

We implemented our seeding method in a short read aligner, strobealign. We demonstrate that strobealign achieves comparable accuracy and percentage of aligned reads to the established aligner BWA-MEM when aligning paired-end reads of several different lengths from several genomes (Figs. 3A, 4A, Suppl. Fig. S2, S12, and S16) while being 6-9 times faster on most benchmarked genomes for read lengths 150nt and longer (Figs. 3C, 4C, Suppl. Fig. S3 and S13). Strobealign is also typically 3.5-7 times faster and has 2-2.5 times lower memory usage than BWA-MEM2 (Suppl. Fig. S14). Notably, we also observe that strobealign is both more accurate and several times faster than BWA-MEM and BWA-MEM2 on some genomes (SIM4, drosophila, REPEATS) for read lengths of 150nt and longer. While other aligners fare well on individual experiments, they do not generally achieve state-of-the-art accuracy on several datasets caused by *e.g*., sequence diversity, repetitive genomes, or longer read lengths. Our conclusions from our simulated experiments also translate to the biological datasets. Strobealign has the most aligned reads (Fig. 5A), the largest concordance with state-of-the-art BWA-MEM and Bowtie2 (Fig. 5B), and the fastest runtime (Fig. 5C, Fig. 6C). In addition, strobealign’s alignments achieve the highest F-scores among all aligners when calling SNPs on the biological data sets and indels on the simulated datasets. Our experiments suggest that for Illumina reads of 150nt and longer, strobealign can remove the alignment bottleneck in many analysis pipelines without compromising mapping accuracy and downstream SNP and indel calling. A caveat with assessing an aligners accuracy based on downstream variant calls is that a caller may take the MAPQ score into account when calling variants, resulting in some variant callers being optimized for the scores of specific aligners. Therefore, it is important to take both mapping accuracy and variant calling results into account in assessing the accuracy of an aligner.

The memory usage of strobealign and other recent aligners (Accel-Align and SNAP) are relatively high compared to memory-efficient tools such as BWA-MEM and Bowie2 (Suppl. Fig. S3). However, most large sequencing datasets are aligned in multi-thread mode on computing resources with many cores and a RAM higher than 32Gb. Therefore, the memory constraint should not be of practical concern in many common bioinformatic pipelines.

### Future work

A large part of the runtime for the BIO150 and BIO250 datasets is spent in base level alignments with ssw (Fig. 2). It may therefore be possible to further optimize runtime by considering a faster Smith-Waterman alignment such as the Wavefront Alignment Algorithm (36) as discussed in (21). Or find a better strategy based on the seeds to select fewer candidate sites to align to. Memory optimizations could also be investigated, such as changing hash values from 64-bit to 32-bit representation (Suppl. Note G). However, these optimizations may come at the cost of accuracy or limitations to the maximum index size.

As for the seeding method, we created randstrobes from syncmers. Other subsampling techniques may further improve the accuracy. While developing strobealign, we tested only minimizers (16) and syncmers (22) and found syncmers to perform better. Word-based methods such as minimally overlapping words (37) have demonstrated to have better conservation than syncmers. However, they are less flexible as syncmers do not need to be pre-computed for different parametrizations. A second direction of exploration is the skewed sampling. During the development of strobealign, we observed that our implemented skewed sampling towards shorter seeds increased accuracy for the shortest reads (≤ 150nt) but had little to no effect compared to uniform sampling for longer reads. We believe it is beneficial for the short reads because more syncmer-pairs will be selected consistently between the read and references near the ends of the read, where there are few syncmers left to sample. It is possible that, e.g., a sampling skew towards longer seeds may be beneficial for longer reads that are not as dependent on single matches and can instead leverage increased uniqueness from longer seeds. Further work on seed length sampling distributions and subsampling densities could be explored.

### Our seeding method in other applications

A natural future research direction is to adapt our seeding method to other mapping scenarios. For example, applications such as long-read alignment (17, 20) or transcriptomic long-read clustering (38) may be substantially sped up when using longer, more unique seeds. Our sampling technique may also improve the computation speed of seeds for transcriptomic spliced alignment (39) as MEM finding is the current bottleneck. Another interesting direction is to explore using our seeding approach for overlap detection for genome assembly. Our seeding method can be thought of as being constructed in syncmer-space instead of over the entire sequence (sequence space). In genome assembly and error correction, ideas to work in minimizer-space instead of in sequence-space have been proposed through the use of paired minimizers (40, 41) and *k* consecutive minimizers (*k*-min-mers) (42) to represent overlaps and assembly graphs.

## Conclusion

We presented a novel strategy to compute seeds based on syncmers and strobemers that can be used for candidate mapping-site detection in sequence mapping applications. We showed that our seeding is fast and used a novel metric E-hits to demonstrate our seeding method’s effectiveness at removing the repetitiveness of seeds. We implemented our seeding strategy in a short-read aligner strobealign. For read lengths of 150nt and longer, strobealign is several times faster than traditional aligners with comparable accuracy. Strobealign can remove the alignment bottleneck in many bioinformatic analysis pipelines and free up substantial computing resources. Furthermore, our seeding approach can potentially be used in many other applications that require sequence mapping.

## Data availability

Strobealign is available at https://github.com/ksahlin/StrobeAlign (v0.4 was used in the benchmark). Script to evaluate seeding methods, generate simulated data, and to evaluate the aligners on all the datasets are available at https://github.com/ksahlin/alignment_evaluation. Biological datasets BIO150 (Illumina WGS 2×150bp, HG004) and BIO250 (Illumina WGS 2×250bp, HG004) analyzed in this study are found at https://github.com/genome-in-a-bottle/giab_data_indexes.

## Methods

### Definitions

We use *i* to index the position in a string *S* and let *S*[*i, k*] denote a *k-mer* substring at position *i* in *S* covering the *k* positions *i*, …, *i* + *k* − 1 in *s*. We will consider 0-indexed strings. We let | · |denote both the length operator applied to strings and the cardinality operator applied to sets. We refer to a *subsequence* of a string as a set of ordered letters that can be derived from a string by deleting some or no letters without changing the order of the remaining letters. A substring is a subsequence where all the letters are consecutive. Our fuzzy seeds produced from *S* are subsequences of *S* since they consist of two syncmers *k*_1_ and *k*_2_ that do not necessarily overlap on *S*. The syncmers are concatenated after their extraction to a string *k*_12_ ≐ *k*_1_*k*_2_ that constitutes the strobemer seed.

We use 2-bit encoding to store nucleotides with 00, 01, 10, and 11 representing A, C, G, and T. With this bit encoding in mind, each string is associated with a sequence of bits that can be interpreted as an integer. For example, *TCA* = 110100 = 52. From now on, we will assume that all strings are manipulated by manipulating their sequence of bits. We use *h* to denote a hash function *h* : {0, 1} ^∗ ⟶^ {0, 1} ^∗^ mapping a sequence of bits to another sequence of bits. Finally, we say that two seeds *k*_12_ and 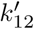 match if 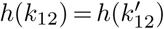 We will denote a match as *m* and use *m.q*_*s*_, *m.q*_*e*_, *m.r*_*s*_, *m.r*_*e*_, *m.o* to denote the read start and end positions, the reference start and end positions, and the orientation of the match, respectively.

### Open syncmers

Open syncmers is a *k*-mer subsampling method described in (22). They are sampled based on three parameters (*k, s, t*). The open-syncmer method compares the *k* − *s* + 1 consecutive *s*-mers within a *k*-mer and selects the *k*-mer as a syncmer if the smallest *s*-mer occurs at position *t* ∈ [0, *k* − *s* + 1] within the *k*-mer. With the smallest *s*-mer, we mean the hash value that the *s*-mer produces. Similar to what is commonly performed in *k*-mer applications, we use a canonical representation of syncmers. A canonical representation means that the lexicographically smallest syncmer out of its forward and reverse-complement sequence is stored.

### Strobemers

Strobemers are described in (23) and consists of several shorter *k*-mers, referred to as *strobes*. We use the randstrobe method (23) described below. Four parameters (*n, k, w*_*min*_, *w*_*max*_) are used to define a randstrobe where *n* is the number of strobes, *k* is the length of the strobes, and *w*_*min*_ and *w*_*max*_ is the lower and upper coordinate offset to the first strobe *k*_1_ for sampling the second strobe *k*_2_ on a string *S*. We will here consider randstrobes with *n* = 2, *i.e*., consists of two strobes. Let *k*_1_ have coordinate *i* on *S* and be the set of *k*-mers from the substring (window) *S*[*i* + *w*_*min*_ : *i* + *w*_*max*_ + *k* − 1]. Then, the randstrobe samples strobe *k*_2_ according to the following sampling function (23):

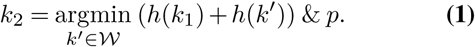

where & is the bitwise AND operator and *p* is a 64-bit bit-mask consisting of 1’s at the *p* leftmost bits and remaining 0’s.

### Modifications to strobemers

Strobemers was first described as being produced over the set of all *k*-mers in a sequence (23). The first modification we make to strobemers as originally described is that we will compute the strobemers over syncmers. Therefore, from now on *k*_1_, *k*_2_, and *k*′ are used to denote the subsampled set of *k*-mers that are open syncmers. We will let [*w*_*min*_, *w*_*max*_] refer to the lower and upper number of syncmers downstream from *k*_1_ where we sample *k*_2_ from. Assume that *k*_1_ has coordinate *i*, then we let W_*s*_ denote the set of syncmers in the substring *S*[*i* + *w*_*min*_ : *i* + *w*_*max*_ + *k* − 1].

A second modification to the strobemers (23) is that we use a skewed sampling function that selects nearby syncmers more frequently. The sampling skew for sampling the second syncmer *k*_2_ is produced from

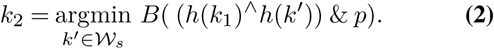

Here, ^∧^ is the bitwise XOR operator, and *B* counts the number of set bits. In other words, *B* returns the number of set bits among the *p ∈* [1, 64] leftmost bits in the 64-bit integer produced after the XOR operation of the hash values of the two strings. Function 2 maps the value space down to [0,p-1] and collisions are resolved by picking the first leftmost strobe. Therefore, a lower value on *p* results in more often picking nearby strobes. An example distribution is shown in Suppl Fig. S1. We found that function 2 gave significantly improved performance on shorter reads (100-150nt) compared to function 1.

The third modification to strobemers is our stored hash value for the strobemer. We store for *k*_12_ the value *h*′ (*k*_12_) ≐ *h*(*k*_1_)/2 + *h*(*k*_2_)/2. The hash function *h*′ is symmetric (*h*′ (*k*_12_) = *h*′ (*k*_21_)) and together with canonical syncmers, it produces the same hash value if the strobemer is created from forward and reverse complement direction. It is stated in (23) that symmetrical hash functions are undesirable for mapping due to unnecessary hash collisions. However, when masking highly repetitive seeds as commonly performed in aligners (18), it turns out that a symmetrical hash function helps to avoid sub-optimal alignments when using strobemers. The reason is the same as for using canonical *k*-mers in read mapping and overlap detection algorithms. Namely, it allows for consistent masking and treatment of forward and reverse complement mapping locations. We will now describe why.

Assume we would use an asymmetric hash function, such as *h*^*″*^ (*k*_12_) = *h*(*k*_1_)/2 + *h*(*k*_2_)/3 proposed in (23). Also assume that strobemer seeds *k*_12_ and *k*_21_ are both found in forward orientation the reference due to, *e.g*., inversions. In this case, only *k*_12_ may be masked because of its distinct hash value to *k*_21_. Now, consider a read in which we extract *k*_12_ in forward direction and *k*_21_ in reverse complement direction. If the read has an optimal match to forward direction with seed *k*_12_ (masked on reference), we would still find the suboptimal match of *k*_21_ of the read in reverse complement orientation to the reference. By using a symmetric hash function, we guarantee to mask the same strobemers in both directions. We observed that such cases are common on, *e.g*., chromosome X in the human genome.

Another benefit of this symmetrical value which does not hold for exact seeds, is that we can use *false symmetrical matches* to our benefit. A false symmetrical match is when the forward seed starting with syncmer *k*_1_ and reverse complement staring with syncmer *k*_2_ become linked as seeds *k*_12_ and *k*_21_, respectively, hence *h*′ (*k*_12_) = *h*′ (*k*_21_). This happens relatively frequently but is not guaranteed. That is, even if the minimizing syncmer for *k*_1_ is *k*_2_, *k*_1_ does not need to be minimizing syncmer *k*_2_. However, the beneficial scenario happens when we have a false symmetric match and, for example, the forward orientation seed is destroyed because of mutations. In this case, it is not guaranteed that the match in the other orientation is destroyed. Thus, we get a false symmetrical match that helps us locate the read location on the genome, which is useful for reads with very few matches. The event of false symmetrical matches was realized and implemented in 0.6.1 in strobealign, leading to slightly improved accuracy.

### Indexing

We first construct open syncmers from the reference sequences and then link two open syncmers together using the randstrobe method with equation 2 as sampling function. A beneficial characteristic with open syncmers is that the the same syncmers will be created from the forward and reverse complement strand if *k* − *s* + 1 is odd and *t* = ⌈(*k* − *s* + 1)/2⌉. Conveniently, it was shown that choosing *t* = ⌈(*k* − *s* + 1)/2⌉ is the optimal parameter for sequence mapping (24). We compute such canonical open syncmers (using *k* = 22, *s* = 18, *t* = 3 as default values) to produce a subsampling of roughly 20% of the *k*-mers sampled, which is similar to *w* = 10 in the minimizer sampling method. As for forming the strobemers, we compute the second strobe from a window of [*w*_*min*_, *w*_*max*_] downstream syncmers to the first strobe, where we set *w*_*min*_ and *w*_*max*_ dependant on the read length based on an experimental evaluation. See details in Section Implementation details.

We store tuples (*h*′ (*k*_12_), *r*_*s*_, *v*) in a flat vector *V* where *h*′ (*k*_12_) is the 64-bit integer hash value of the strobemer, *r*_*s*_ is the coordinate start of the first strobe (32-bit integer), and *v* is a 32-bit integer containing reference id (rightmost 24 bits) and the offset of the second strobe (leftmost 8 bits). We sort *V* by hash values and construct a hash table with hash values as keys pointing to offset and the number of occurrences of the hash value in *V*. By lookup of *h*′ (*k*_12_), we know which segment in *V* to iterate over to find query matches. This type of index representation has been used previously (18, 21), and was suggested to us by Heng Li (43). Finally, similarly to minimap2, we mask (ignore) a top fraction of repetitive strobemers in the reference. This value is a parameter to strobealign and is by default set to 0.0002, similarly to minimap2.

### Finding candidate mapping sites

Strobealign computes canonical open syncmers from the read similarly to what is described above to index the reference. Since the created syncmers are canonical, we can compute forward and reverse complemented strobemers by iterating over the syncmers forward and reverse order, respectively. Computing strobemers in forward and reverse orientation gives us a vector of tuples (*q*_*s*_, *q*_*e*_, *o*) representing start coordinate *q*_*s*_, end coordinate *q*_*e*_, and orientation *o* (Boolean value with 0 representing forward and 1 representing reverse complement), of the strobemer on the read.

If a strobemer is found in the reference, it will have one or more coordinate-tuples (*r*_*id*_, *r*_*s*_, *r*_*e*_) in the reference. We call *m* = (*r*_*id*_, *r*_*s*_, *r*_*e*_, *q*_*s*_, *q*_*e*_, *o*) a *match* between the query and the reference. Let *d* be the difference in length between the strobemer on the reference and query. If several matches are found, Strobealign iterates over the matches, stores the lowest observed *d* during iteration, and saves only the matches with the current lowest *d*. This approach is not the same as first computing the lowest *d* and iterating a second pass to store only matches with the lowest *d*. We chose the former as we tried both and observed close to no difference in accuracy while slightly increasing runtime in the latter case due to the extra iteration.

From the stored matches, strobealign constructs *merged matchesℳ*which are similar but slightly more stringent to Non-overlapping Approximate Matches (NAMs) that are defined in (23). Merged matches are produced as follows. We iterate over all matches in ascending order in the read and join two matches *m* and *m*^*′*^ into a merged match if it holds that

i. (*m.r*_*id*_ == *m*^*′*^ .*r*_*id*_) && (*m.o* == *m*^*′*^ .*o*)
ii. *m.q*_*s*_ *< m*^*′*^ .*q*_*s*_ ≤ *m.q*_*e*_
iii. *m.r*_*s*_ *< m*^*′*^ .*r*_*s*_ ≤ *m.r*_*e*_
iv. and if one of the following holds; (*m.q*_*s*_ *< m*′ .*q*_*s*_ *< m.q*_*e*_ *< m*′ .*q*_*e*_) ∧ (*m.r*_*s*_ *< m*′ .*r*_*s*_ *< m.r*_*e*_ *< m*′ .*r*_*e*_) or (*m.q*_*s*_ *< m*′ .*q*_*s*_ *< m*′ .*q*_*e*_ *< m.q*_*e*_) ∧ (*m.r*_*s*_ *< m*′ .*r*_*s*_ *< m*′ .*r*_*e*_ *< m.r*_*e*_).

In other words, the matches need to (i) come from the same reference and have the same direction, (ii-iii) overlap on both query and reference, and (iv) two strobemer matches need to have a consistent ordering of the four strobes on the reference and the query. Specifically, a NAM requires only (i-iii). There are scenarios due to local repeats where, for example, (*m.q*_*s*_ *< m*′ .*q*_*s*_ *< m.q*_*e*_ *< m*′ .*q*_*e*_) is the order on the query but (*m.r*_*s*_ *< m*′ .*r*_*s*_ *< m*′ .*r*_*e*_ *< m.r*_*e*_) is the order on the reference invalidating (iv). We consider such cases as separate matches.

If *m* is the current considered match in the iteration over the matches in a query, we refer to all matches *m* ^*″*^ with *m*^*″*^ .*q*_*s*_ *< m.q*_*s*_ ≤ *m*^*″*^ .*q*_*e*_ as *open matches* and *m*^*″*^ .*q*_*e*_ *< m.q*_*s*_ as *closed matches*. While iterating over the matches in a read, we keep a vector of currently open *merged matches*, and filter out the closed matches in this vector. In a merged match ℳ we keep information of how many matches were added, the position of the first and last strobe on query and reference, and the orientation on the reference genome. After the final match on the query, we close all merged matches. The closed matches are the final merged matches that constitute the candidate mapping locations.

### Computing MAPQ score

After merging matches, each merged match ℳ consists of a number of matches | ℳ | and a total span-range of the merged match on both the query *a* = ℳ.*q*_*e*_ − ℳ.*q*_*s*_ and the reference *b* = ℳ.*r*_*e*_ − ℳ.*r*_*s*_. We define the score of ℳ as *S*_ℳ_ = (min {*a, b*} − |*a* − *b*|)· | ℳ |, which acknowledges only the minimum span over the query and reference and penalizes if there is a difference in the span lengths. We compute the MAPQ score similarly to minimap2 but substitute minimap2’s chain score (18) to our merged match score *S*_ℳ_. That is, if *S*_1_ and *S*_2_ are the top two scoring merged matches for a read, the MAPQ is computed by

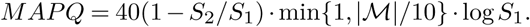

### Single-end Alignment

Merged matches are produced and scored as described above and constitute the candidate mapping regions. For each candidate region sorted order with respect to the score, we extract segments on the reference defined by coordinates (ℳ.*r*_*s*_ − ℳ.*q*_*e*_, ℳ.*r*_*e*_ + (|*q*| − ℳ.*q*_*e*_)), where |*q*| is the length of the read. If | ℳ .*q*_*e*_ − ℳ.*q*_*s*_| = | ℳ.*r*_*e*_ − ℳ.*r*_*s*_ |, we compute the Hamming distance between the read and the extracted reference segment. Otherwise, if the distance between merged match is different on the reference and query due to, *e.g*., indels, we send the sequences to alignment with ssw (44). We use a match score of 1 and alignment penalties of 4, 6, and 1 for mismatch, gap open, and gap extension, respectively. Additionally, if the computed Hamming distance is larger than 0.05 |*r*| where |*r*| denotes the read length, we perform an additional alignment with ssw as, theoretically, there may be more than one indel within the mapped location that would lead to the same match lengths on the read and the reference.

#### Rescue mode

A read could have few or zero matches if all the strobemers extracted from the read were masked due to being too abundant. The abundance cutoff, which we denote as *A*, is controlled with a parameter -f (default value 0.0002), as in minimap2. For example, *A* is between 30 and 50 for hg38 depending on the values we use for parameters *k, w*_*min*_, and *w*_*max*_, described in Section Implementation details. If more than 30% or strobemers were masked when finding matches, strobealign enters a rescue mode where it considers a higher threshold. In the rescue mode, strobealign sorts the seeds according to the abundance on the reference. Then it selects all the seeds below an abundance of *R* (*R* is a positive integer parameter with default value 2) and, if this still produces fewer than 5 seeds, it uses 1000 as a hard abundance threshold. If there are still 0 matches, the read is treated as unmapped.

### Paired-end Alignment

Similar to the single-end mapping mode, strobealign computes merged matches for both mates within the read pair and employs an identical rescue mode if there are too many masked strobemer seeds, as described for the single-end mapping. There are, however, two additional components in the paired-end mapping mode. Firstly, strobealign employs a joint scoring of candidate mapping location based on expected insert size (similar to BWA-MEM). Secondly, based on the mate’s mapping location, strobealign can enter a rescue mode even for a read with zero matches. We describe the two components below.

For the joint scoring, strobealign first sorts the candidate map locations based on the total seed count for the two mates in a read pair. Then, strobealign finds the best candidate locations from a combined MAPQ score described below. Let 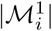 and 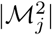 be the number of matches in i-th and j-th merged match for the first and the second mate in the read-pair, respectively. If it holds for some *i* and *j* that the two merged matches are on the same chromosome in the correct relative orientation with a mapped distance *< μ* + 10*σ*, the joint map-location count *C*_*ij*_ is simply 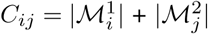. We also add the individual candidate map location counts obtained as 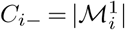 and 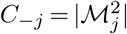 for the two mates individually. For the scores in order of highest total seed count first, strobealign performs base-level alignment of each mate (as described in the Single-end alignment section). The alignment score *S*_*ij*_ of such aligned pair is then computed as

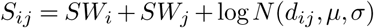

where *SW* denotes the Smith-Waterman alignment score, and *d*_*ij*_ denotes the distance between the mates on the genome. The individually mapped reads (*e.g*., if on different chromosomes) are given a score

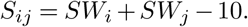

Since -10 corresponds to more than 4 standard deviations away, this is the score cutoff which prefers the reads to be mapped individually at their respective locations with the highest SW score.

#### Mate rescue mode

If one of the mates does not have any merged match, we perform base level alignment of the mate without merged match within a genomic segment of [0, *μ* + 5*σ*] nucleotides away in the expected direction from the location of the mate with a merged match.

### Implementation details

Similar to the default parameters in minimap2, we consider the top 20 MAPQ (or joint MAPQ) scoring candidates for alignment, and we implement a dropoff score threshold of 0.5 (score to the highest score). In addition to these parameters, we employ two additional optimizations. First, if we encounter a perfect match (no mismatches), we stop and report the alignment even if there are remaining candidates above the drop-off parameter. Second, suppose we have encountered an alignment with an edit distance of 1. In that case, we do not call base-level alignment for remaining candidates since a call to base level alignment implies that the edit distance is at least 1, as we described in the single-end alignment section. Strobealign also supports multithreading using openMP. If more than one thread is specified, strobealign will parallelize the alignment step by splitting the reads into batches of 1 million reads to be processed in parallel.

The median read length 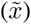 is estimated from the first 500 reads in the read input. As for selecting values of *k, s, p, w*_*min*_ and *w*_*max*_, we set suitable parameters given 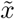 based on experimental evaluation of accuracy and runtime. We let *w*_*min*_ = *k/*(*k* − *s* + 1) + *l* and *w*_*min*_ = *k/*(*k* − *s* + 1) + *u* where *l* and *u* are integers that specify lower and upper offset. Then we choose the following parameter tuples for (*k, s, p, l, u*) given 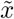.

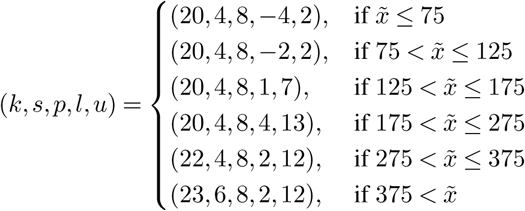

With a read length of length 200nt and the parameters in this study, roughly (1/5) ∗ 200 = 40 syncmers are produced for each read, and roughly 30 strobemers are created in each direction. Naturally, these values are reduced for shorter reads which impacts sensitivity. A read of 100nt will have on average 20 syncmers and only 10 strobemers. We could consider lower *w*_*max*_ to produce more strobemers for shorter reads at the cost of memory.

Although it is exponentially less likely to have open syncmers sampled further away from the mean sampling density (22), they do not have a window guarantee and may be sparser sampled in some regions. Therefore, we have a hard limit on the maximum seed size as a parameter to strobealign (*m*), where it defaults to 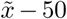. This means that, in some cases, the maximum seed size may be lower than the downstream nucleotide level distance of the syncmer corresponding to *w*_*max*_. With the parameters we use above, this happens in less than 0.1% of the seeds. Finally, on rare occasions there is no open syncmer within the downstream window, *e.g*., due to regions of N’s in centromeres on hg38. In these cases we use only the first syncmer as the strobemer seed. This happens on hg38 for 0.0007% of the 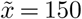 seeds (3,963 out of 544M seeds).

### The E-hits metric

Let *N* be the total number of seeds and *M* ≤ *N* the number of distinct seeds produced over a set of reference sequences by any seeding method. Let *i ∈* [1,*M*] be an index variable over the set of distinct seeds, and *x*_*i*_ denote the number of times seed *i* is produced. Then, 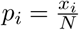 corresponds to the probability that seed *i* is picked uniformly at random over the multiset of seeds produced by the seeding method. We let our set of observations *x*_*i*_ ∈ 𝒳 and our probability distribution *p*_*i*_ ∈ 𝓅 be the model of the scenario of sampling uniformly at random a location on the reference sequences and extracting a seed. Then, the expected number of occurrences of a randomly sampled seed *i* on the reference is

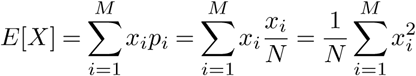

We refer to *E*[*X*] as E-hits. The connection to read mapping is immediate. If reads are sampled uniformly at random over the reference sequences, E-hits measure the expected number of matches we obtain for error-free seeds extracted from the reads. The uniform distribution is a common assumption for, e.g., Illumina genome sequencing reads, albeit not fully accurate. The E-hits metric is conceptually similar to the expected contig size covering a random position in the genome (E-size) as defined in (45), hence E-hits’ similar denotation. Note that E-hist is different from the popular E-value used in BLAST (46). The E-value is a theoretical computation that measures the expected number hits that could be found by chance under random nucleotide distribution given a database size and an exact *k*-mer seed. E-hits is computed from the actual reference sequences and any seeding protocol.

### Memory usage

Strobealign has a peak memory usage of about 25-33Gb for hg38 (Suppl. Fig. S5). With the parameter settings we investigated in this evaluation for read lengths 50-300nt on hg38 (syncmer subsampling rate of 1/5), strobealign stores roughly 544 million seeds in memory. Furthermore, the size of the index scales with the number of unique seeds. For example, strobealign uses only about 1.5 (50Gb) more memory when aligning to rye (2.3 times hg38 in size) as rye is a repetitive genome with a lower fraction of unique seeds. Similarly, the maize index is only 15Gb (0.5 times the index of human). A detailed discussion on the memory and implementation is found in Suppl Note G.

## ACKNOWLEDGEMENTS

The computations were performed on resources provided by the Swedish National Infrastructure for Computing (SNIC) at Uppsala Multidisciplinary Center for Advanced Computational Science (UPPMAX) partially funded by the Swedish Research Council through grant agreement no. 2018-05973.

