## supplemental_materials for "Flexible seed size enables ultra-fast and accurate read alignment"

Kristoffer Sahlin

Department of mathematics, Science for Life Laboratory, Stockholm University, 106 91, Stockholm, Sweden

#### Note A: Simulations

##### hg38 simulations

We used mason2 [2] to simulate VCF files with variants to hg38 at various SNP and indel rates, and denote the datasets SIM1, SIM2, SIM3, and SIM4 for increasing degree of mutation and indel rates to hg38. For SIM1-SIM4 we set the following parameters to mason_variator

SIM1: –sv-indel-rate 0.0000001 –snp-rate 0.0001 –small-indel-rate 0.000001 –max-small-indel-size 6 (default mason_variator parameters)

SIM2: –sv-indel-rate 0.000001 –snp-rate 0.001 –small-indel-rate 0.00001 –max-small-indel-size 20

SIM3: –sv-indel-rate 0.000005 –snp-rate 0.001 –small-indel-rate 0.0001 –max-small-indel-size 50

SIM4: –sv-indel-rate 0.00001 –snp-rate 0.005 –small-indel-rate 0.0005 –max-small-indel-size 50

For each of the genomes SIM1, SIM2, SIM3, and SIM4, we used mason_simulator to simulate Illumina single and paired-end reads with read lengths 50, 75, 100, 150, 200, 250, 300, and 500. For the 250, 300, and 500 paired-end read experiments, we had to set the parameter –fragment-mean-size 700 (default is 300) as mason_simulator could not complete due to he fragment size being too small for the read length. We then mapped the reads simulated from SIM1, SIM2, and SIM3 to hg38 with the aligners.

#### REPEATS simulations

We simulated a string of 100,000nt by choosing letters A, C, G, and T at random. We produced 500 copies of this string but introduced a 5% SNPs and deleted segments of length between 1 and 1000 with probability 0.0001 on each copy. This roughly represents 500 copies of length 90-100kbp at a rough 90% identity between copies with some deletions of various sizes and locations.

We furthermore simulated reads from a related genome to the above repetitive genome using mason_variator with the parameters –sv-indel-rate 0.00005 –snp-rate 0.005 –small-indel-rate 0.0005 –max-small-indel-size 50. We denote this dataset REPEATS.

#### Drosophila, maize, CHM13, and rye simulations

We used mason2 to simulate VCF files with variants to each respective genome at the same SNP and indel rate as SIM3 for human, and denote the datasets drosophila, maize, CHM13, and rye. Reads were simulated in the same way as for the experiments on hg38.

For drosophila we used assembly BDGP6.22 (available here: `ftp://ftp.ensembl.org/pub/release-97/fasta/drosophila_melanogaster/dna/Drosophila_melanogaster.BDGP6.22.dna.toplevel.fa.gz.`). For maize we used the ZM version (available here: `https://download.maizegdb.org/Zm-B73-REFERENCE-NAM-5.0/Zm-B73-REFERENCE-NAM-5.0.fa.gz`). For CHM13, we used assembly v2.2 (available here: `https://s3-us-west-2.amazonaws.com/human-pangenomics/T2T/CHM13/assemblies/analysis_set/chm13v2.0.fa.gz`). For rye genome, we used assembly GCA_902687465.1 (available here: `https://ftp.ncbi.nlm.nih.gov/genomes/genbank/plant/Secale_cereale/latest_assembly_versions/GCA_016097815.1_HAU_Weining_v1.0/GCA_016097815.1_HAU_Weining_v1.0_genomic.fna.gz`).

##### Excluded rye contigs

Mason2 opens temporary files for each contig when simulating reads. The rye genome contains about 25,000 contigs. Due to a contraint on the maximum number of files open at the same time on our system we has to remove some contigs in order to simulate reads. We therefore reduced the number of contigs by removing all contigs shorter than 50,000nt. This resulted in 528 remaining contigs and a total reference size of about 7.3 billion nucleotides.

##### Excluded Drosophila contigs

We excluded two contigs that were smaller than 700nt in the drosophila genome because for the longer insert size simulations, mason_simulator could not sample a fragment smaller than the contig size from these contigs and returned an error.

### Note B: Evaluation metrics

We measured the accuracy, percentage of aligned reads, runtime, peak memory usage, and over-aligned reads for each of the tools.

accuracy: The fraction of correctly aligned reads out of all reads. We consider a read correctly aligned if the reported alignment coordinates overlaps with the provided ground truth reference coordinates for the read. Reversely, an incorrectly aligned read is a read that is aligned to a location that does not overlap the true alignment location, i.e., they have disjoint alignment coordinates.

% aligned reads: The percentage of aligned reads out of the total number of reads.

runtime: Time of alignment excluding the time to construct or read the index.

peak memory: Maximum memory footprint obtained by /usr/bin/time -v

Over-aligned: Number of reads aligned by the aligners that are reported as unaligned in the mason ground truth file.

Alignment runtime is measured after the index has been loaded into memory. We can parse and subtract the time to read or construct the index from the standard output of strobealign, minimap2, Accel-Align, and SNAP. As described in [6], larger indexes take a longer time to load from disc, although the difference is negligible when aligning a large set of reads. On our machine, loading the index took about 22 seconds for minimap2, 25 seconds for Accel-Align, 22 seconds for SNAP, and 18 seconds for URMAP. For BWA-MEM and Bowtie2, we did not subtract this time as we cannot parse them from the output, but their indices are smaller and very fast to load (about 6-9 seconds reported in [6]).

As for the over-aligned reads, all the tools aligned an identical number of over-aligned reads meaning that they find locations for these reads (Fig S6, S11). We reported this metric despite not offering any differentiation between aligners.

### Note C: Aligner information

We used default parameters for all the aligners across simulated and biological datasets except for minimap2 and Bowtie2. For minimap2 we used the parameter ”-x sr” for short-read mode. For Bowtie2, we set maximum fragment length -X 1000 (default 500) for the paired-end 250nt and 300nt experiments that we simulated with larger fragment size after communication with Ben Langmead. In addition, we ran Bowtie2 in both default (global alignment) and –local (local alignment) modes. We observed a significant increase in alignment time using local alignment mode. However, we did not observe a noticeable increase in accuracy on simulated data, while it resulted in significantly more reads aligned for the biological datasets. We, therefore, present results for Bowtie2 using the default alignment mode on the simulated datasets, while we present the local alignment results for the biological datasets.

#### Excluded aligners

puffaligner: We could not install puffalignmer on the cluster where we perform our benchmarks, as reported [4].

URMAP: URMAP segfaulted on all 2*300 paired-end experiments, as reported here [3] and could not run on the rye genome as it exited with an error that genome was too big. We also could not run URMAP in multithread mode as reported in the same issue [3]. Since our experiments are all based on running multithreaded mode with 16 threads, we have excluded it from the evaluation. However, we benchmarked URMAP in an earlier version of the study using one thread and found it to be very fast but had a low accuracy and low percentage of aligned reads [5].

#### Excluded individual experiments

SNAP: We excluded SNAP from experiments with read size of 500 as it exited with an error stating that maximum read size is 400. We also excluded SNAP from experiments on the rye genome as it exited with an error message stating that it could not parse a sequence (chromosome) longer than $2^{31}$ bases.

AccelAlign: AccelAlign segfaulted (signal 11) on all of the eight the rye genome datasets and were therefore excluded from experiments on this genome.

### Note D: Biological data

We downloaded the first read pair files containing paired-end read datasets with read lengths 150 and 250 PE from Genome-IN-A-Bottle consortium [7] found in the GitHub repository at `https://github.com/genome-in-a-bottle/giab_data_indexes`. We denote them BIO150 and BIO250. For the BIO150 dataset, all the files with project accession 140818_D00360_0047_BHA66FADXX from HG004 were used. For the BIO250 dataset, all files from HG004 were used. The BIO150 and BIO250 contain 263,877,609 and 104,215,522 million read-pairs which corresponds to a coverage of around 26x and 17x, respectively.

For the mapping concordance analysis, we used a subset of 4 million read-pairs that were taken from the first file in each of the data repositories. The BIO150 reads have file accessions 4A1_CGATGT_L001_R1_001 and 4A1_CGATGT_L001_R2_001, and the BIO250 PE reads have file accessions D3_S1_L001_R1_001 and D3_S1_L001_R2_001. We measured alignment concordance identically to how we computed accuracy for the simulated datasets. That is, the measure is based on an overlap in alignment coordinates.

### Note E: SNP and indel analysis

We use hg38 without alternative haplotypes as proposed in [1] found here (`ftp://ftp.ncbi.nlm.nih.gov/genomes/all/GCA/000/001/405/GCA_000001405.15_GRCh38/seqs_for_alignment_pipelines.ucsc_ids/GCA_000001405.15_GRCh38_no_alt_analysis_set.fna.gz`).

#### Biological variants

We obtain the true SNVs and INDELs from the GIAB gold standard predictions formed from several sequencing technologies. They are provided at `https://ftp-trace.ncbi.nlm.nih.gov/giab/ftp/release/AshkenazimTrio/HG004_NA24143_mother/latest/GRCh38/HG004_GRCh38_1_22_v4.2.1_benchmark.vcf.gz`.

#### Simulated variants

Variants are simulated with mason_variator as for the SIM3 datasets described in Suppl. Note A.

#### SNP and indel calling pipeline

We use *bcftools* to call variants from each of the aligner's output. We split the VCF files with true and predicted variants into separate files. Then we compute the intersection between called and true variations using *bcftools isec* with default options –collapse none, which means that only records with identical REF and ALT alleles are compatible, and are therefore classified as true predictions. We provide pseudocode of these steps below.

```
bcftools mpileup -O z --fasta-ref ref aligned.bam > aligned.vcf.gz
bcftools call -v -c -O v aligned.vcf.gz > aligned.variants.vcf.gz

# Split into SNP and INDELS
grep -v -E -e "INDEL;" aligned.variants.vcf.gz > aligned.variants.SNV.vcf
grep "#"  aligned.variants.vcf.gz > aligned.variants.INDEL.vcf
```

```
grep -E -e "INDEL;" aligned.variants.vcf.gz >> aligned.variants.INDEL.vcf

# Separate GIAB SNVs and INDELS
shell('zgrep "#" true.variants.vcf > true.variants.SNV.vcf')
shell('zgrep -P  "\t[ACGT]\t[ACGT]\t" true.variants.vcf >> true.variants.SNV.vcf')
shell('zgrep -v -P  "\t[ACGT]\t[ACGT]\t" true.variants.vcf > true.variants.INDEL.vcf')

for type in SNV INDEL
do
bcftools sort -Oz aligned.variants.$type.vcf.gz \
               -o aligned.variants.sorted.$type.vcf.gz
bcftools index aligned.variants.sorted.$type.vcf.gz
bcftools isec --nfiles 2 -O u true_variants.sorted.$type.vcf.gz \
              aligned.variants.sorted.$type.vcf -p out_$type
done
```

### Note F: Single-end evaluation

We observe that strobealign is the fastest tool (Fig. S8) for read lengths of 150nt and larger, and SNAP is the fastest for the 100nt reads in our simulated experiments. Strobealign is about 3 times faster than minimap2 and 8 times faster than BWA-MEM across all datasets except the 100nt reads. Furthermore, in comparison to Accel-Align and SNAP, the speedup that strobealign achieves over state-of-the-art aligners does not compromise as much on alignment accuracy (Fig. S9) or deteriorate as quickly with higher diversity (SIM3 and SIM4 datasets). BWA-MEM has the highest accuracy, while minimap2 and strobealign have similar accuracy on the SIM1-SIM3 single-end datasets. Minimap2 achieves a good accuracy-speed tradeoff for the SIM4 dataset.

As for the percentage of aligned reads, they are similar for most tools except for two notable exceptions; (1) SNAP's percent of aligned reads decrease with variation density and read length, and (2) Bowtie2's alignment rate decrease with variation density.

The number of reads reported as over-aligned in our analysis was identical for all tools in alignment mode (Fig. S11), indicating that the tools find places to align the reads tagged as unaligned by mason_simulator.

In summary, strobealign's trade-off between speed and alignment accuracy to the other aligners for the single-end reads lengths of 150nt and above is not as good as for the paired-end experiments. For example, strobealign does not have both highest accuracy and fastest runtime on the high diversity datasets (SIM4) in single-end mode. Still, strobealign offers a very fast alignment time without significant compromise in accuracy.

### Note G: Memory usage during alignment

With the parameter settings we investigated in this evaluation for read lengths 50-300nt on hg38 (syncmer subsampling rate of 1/5), strobealign stores roughly 544 million strobemer tuples in the flat vector. This corresponds to roughly 12Gb (585,809,159*(8+4+4)) for storing the hash value, the start position of the strobe, and a bit-packed 4-byte integer storing both reference id (24 bits) and the second strobe offset to the start strobe (8 bits). The 8-bit allocation for strobe offset puts an upper limit of 255+$k$ on the seed size. The hash table for storing k-mer positions in the flat vector consists of 542,925,043 unique hash values (8 Byte) that points to an offset in the flat vector (4 Byte) and a count (4 Byte). In total, the hash table amounts to roughly 8.7Gb (542,925,043*(8+4+4)) to store only the leaves in the hash table. However, there is a large overhead in storing the internal nodes in the hash map (robin hood hash map in C++). We believe that with additional engineering, we may reduce memory further. Strobealign should only require extra memory for storing the second strobe offset compared to minimap2, as they have a similar index construction. Further, minimap2 and strobealign have similar sampling densities. Minimap2 has an average minimizer sampling density of 2/11 for short reads, while strobealign has a strobemer sampling density of 1/5.

Strobealign does not currently implement a separate indexing step. However, the indexing is fast and the index needs to be rebuilt with different alignment parameters on $k$, $w_{min}$, and $w_{max}$, which makes storing an index much less important compared to BWT based aligners. Nevertheless, separating alignment and indexing could reduce peak memory during alignment by not saving the 64-bit hash value in the flat vector after having built the hash table. However, we believe that most large short-read datasets are usually aligned in multi-threaded mode on computing nodes or computers with more than 32Gb of RAM, for which this memory usage should not be of practical limitation. We outline three possible optimizations to strobealings memory usage:

1. Removing hash values in flat vector after indexing (lower peak for alignment but the same at indexing step)
2. Changing hash values from 64bit to 32bit
3. Using a 16-bit integer to represent count of a seed in the flat vector instead of a 32-bit integer.

Points 1-3 could reduce peak memory substantially, but particularly point2 may impact accuracy and speed.

### Bibliography

[5] Kristoffer Sahlin. Flexible seed size enables ultra-fast and accurate read alignment. *bioRxiv*, 2022.

[6] Yiqing Yan, Nimisha Chaturvedi, and Raja Appuswamy. Accel-align: a fast sequence mapper and aligner based on the seed–embed–extend method. *BMC Bioinformatics*, 22(1):257, 2021.

[7] Justin M. Zook, David Catoe, Jennifer McDaniel, Lindsay Vang, Noah Spies, Arend Sidow, Ziming Weng, Yuling Liu, Christopher E. Mason, Noah Alexander, Elizabeth Henaff, Alexa B. R. McIntyre, Dhruva Chandramohan, Feng Chen, Erich Jaeger, Ali Moshrefi, Khoa Pham, William Stedman, Tiffany Liang, Michael Saghbini, Zeljko Dzakula, Alex Hastie, Han Cao, Gintaras Deikus, Eric Schadt, Robert Sebra, Ali Bashir, Rebecca M. Truty, Christopher C. Chang, Natali Gulbahce, Keyan Zhao, Srinka Ghosh, Fiona Hyland, Yutao Fu, Mark Chaisson, Chunlin Xiao, Jonathan Trow, Stephen T. Sherry, Alexander W. Zaranek, Madeleine Ball, Jason Bobe, Preston Estep, George M. Church, Patrick Marks, Sofia Kyriazopoulou-Panagiotopoulou, Grace X. Y. Zheng, Michael Schnall-Levin, Heather S. Ordonez, Patrice A. Mudivarti, Kristina Giorda, Ying Sheng, Karoline Bjarnesdatter Rypdal, and Marc Salit. Extensive sequencing of seven human genomes to characterize benchmark reference materials. *Scientific Data*, 3(1):160025, 2016.

### Figures

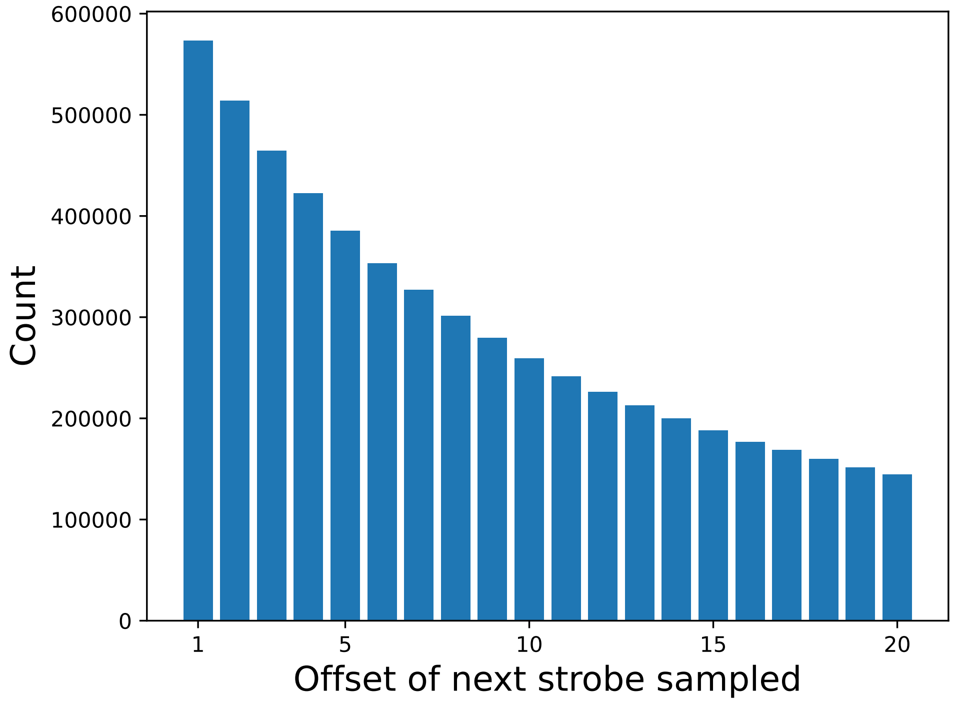


Figure S1: Illustration of the sampling skew of function $B$ using $p = 8$. The figure shows the number of skipped strobes between the first and the second strobe if using $w_{min} = 1$ and $w_{max} = 20$ on a randomly generated DNA string.

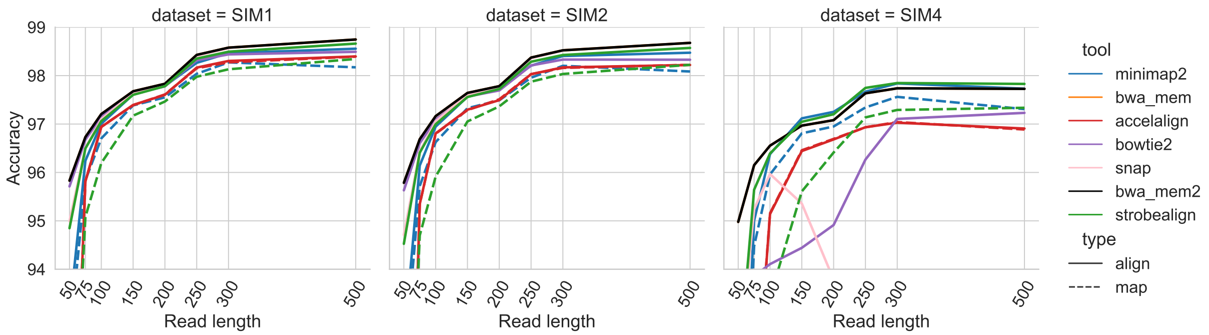


Figure S2: Accuracy of paired-end aligned reads for the SIM1, SIM2, and SIM4 datasets (panels). The x-axis shows simulated read length and the y-axis shows the accuracy.

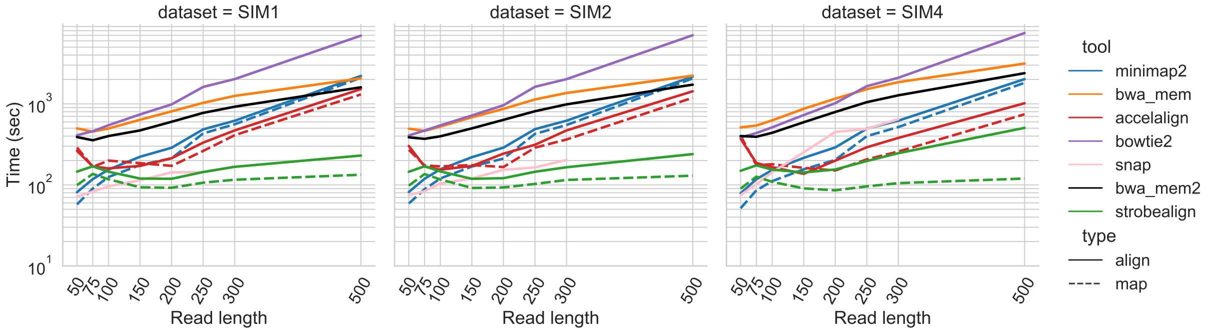


Figure S3: Time to align 10M simulated Illumina paired-end reads for the SIM1, SIM2, and SIM4 datasets (panels). The x-axis shows simulated read length and the y-axis shows the wall clock time from after the index has been loaded into RAM.

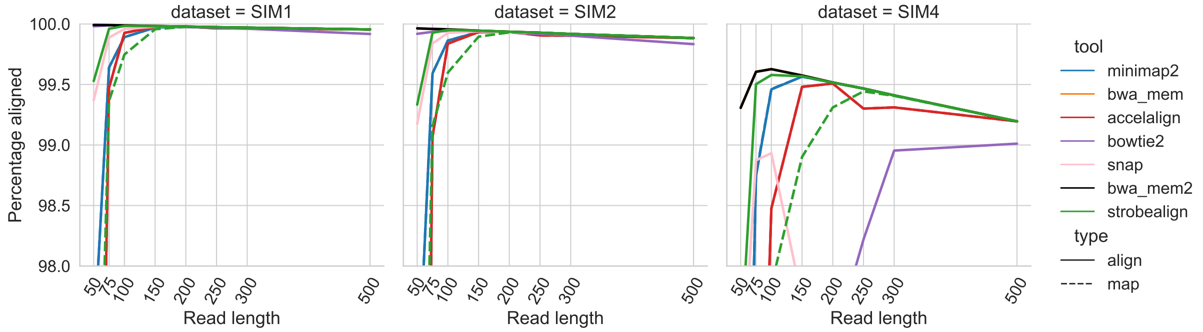


Figure S4: Percent aligned paired-end reads for the SIM1, SIM2, and SIM4 datasets (panels). The x-axis shows simulated read length and the y-axis shows the percentage of aligned reads.

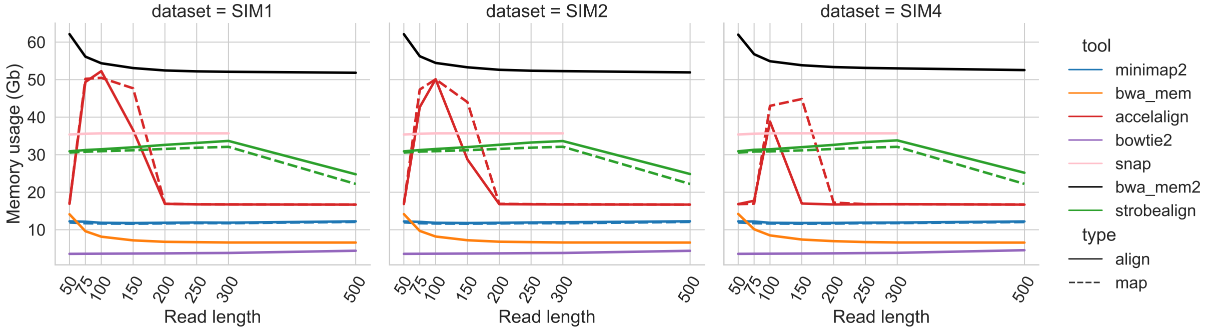


Figure S5: Memory usage across hg38 datasets.

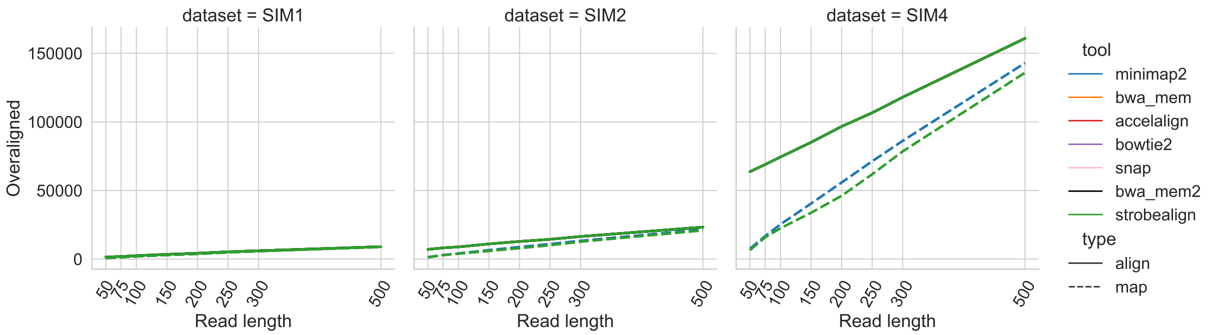


Figure S6: The number of aligned paired-end reads for reads classified as unaligned by the mason simulator. All the aligners have the exact same number of over-aligned read which give plots with only one visible line.

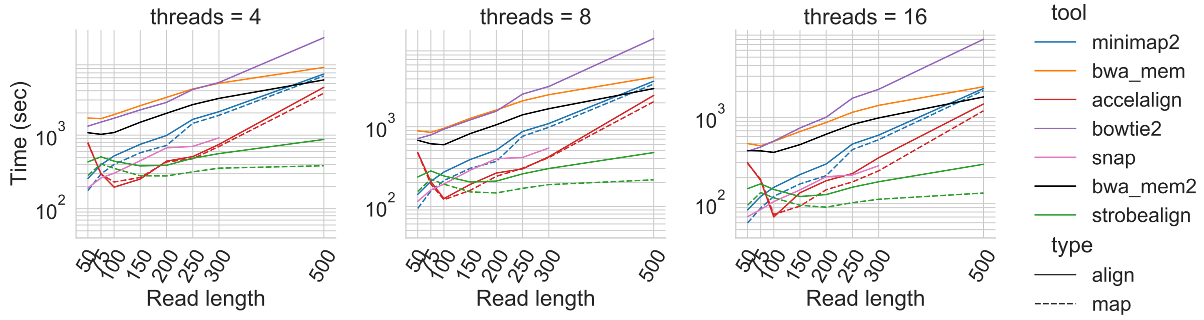


Figure S7: Runtime of aligners with variable number of threads for the paired-end SIM3 datasets.

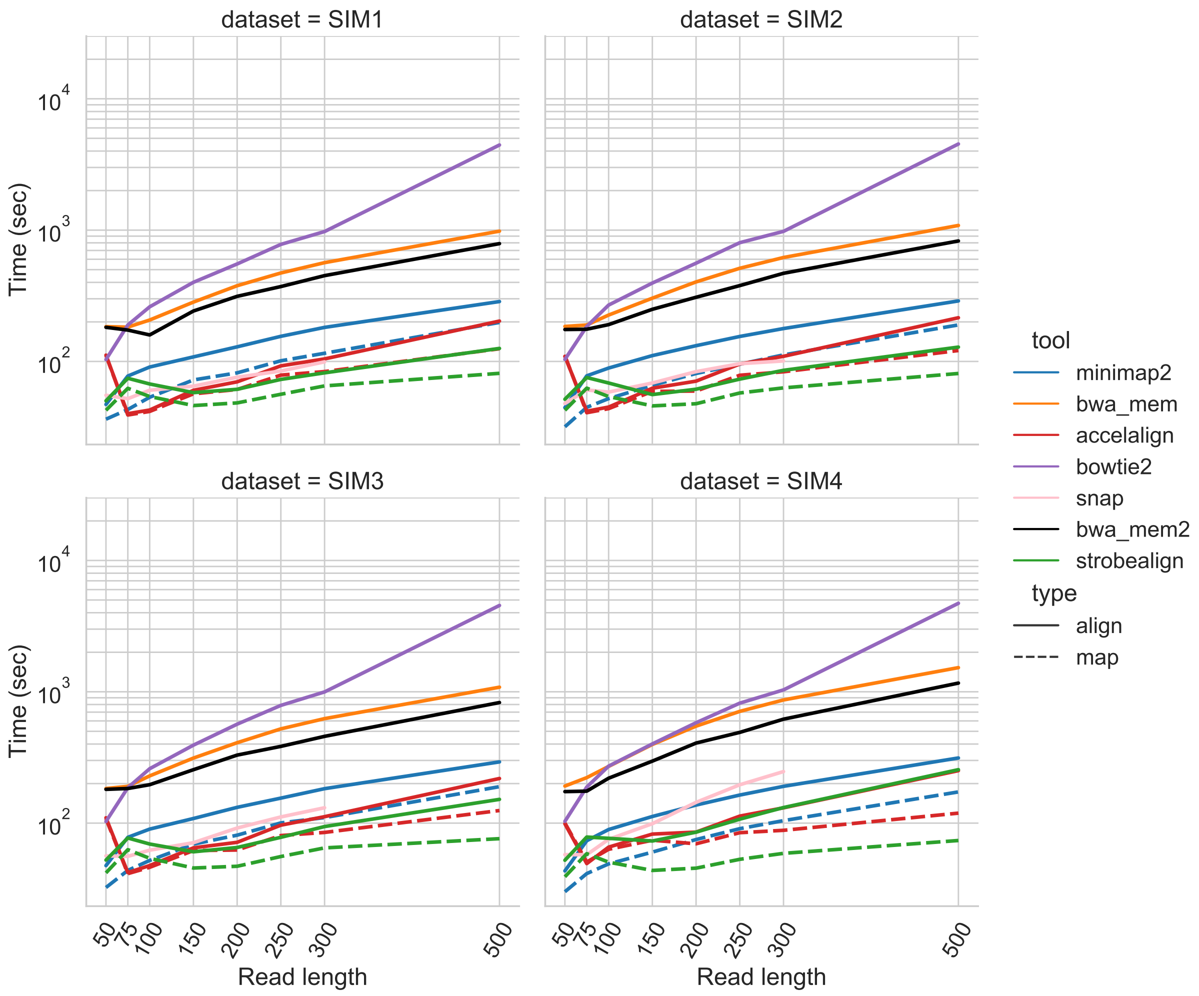


Figure S8: Time taken to align 10M simulated single-end Illumina reads for the three simulated datasets (panels). The x-axis shows simulated read length and the y-axis shows the wall clock time from after the index has been loaded into RAM.

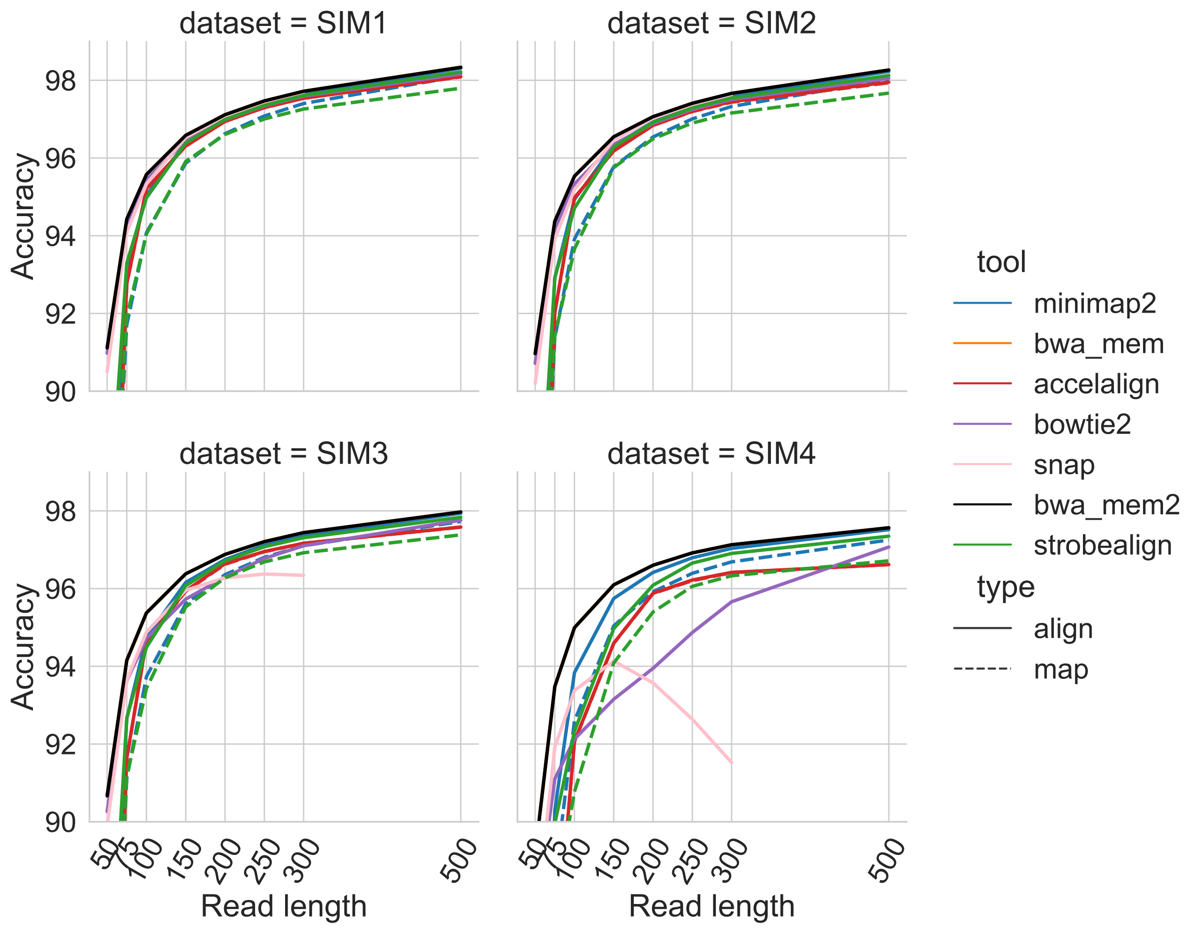


Figure S9: Accuracy of aligned reads for the four single-end Illumina simulated datasets (panels). The x-axis shows simulated read length and the y-axis shows the accuracy.

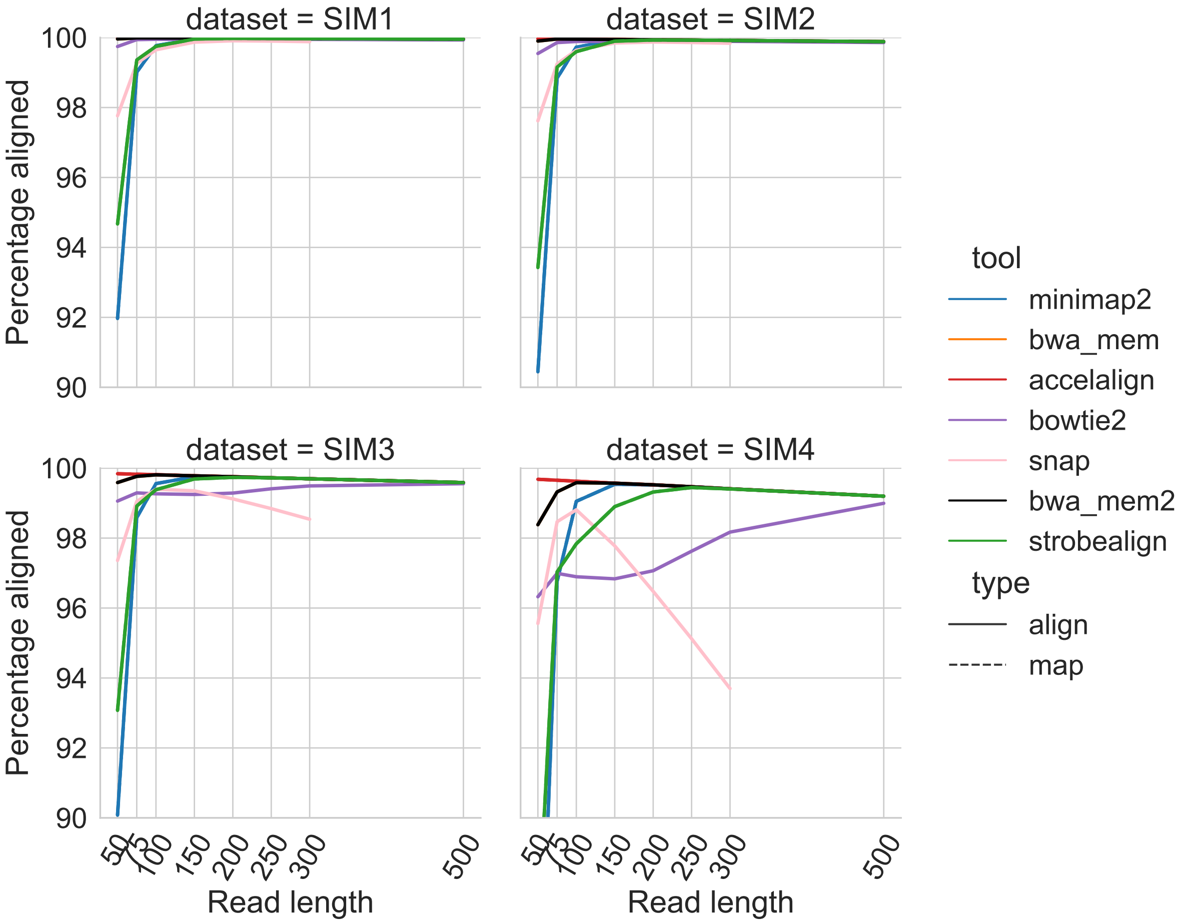


Figure S10: Percent aligned single-end reads for the four single-end Illumina simulated datasets (panels). The x-axis shows simulated read length and the y-axis shows the percentage of aligned reads.

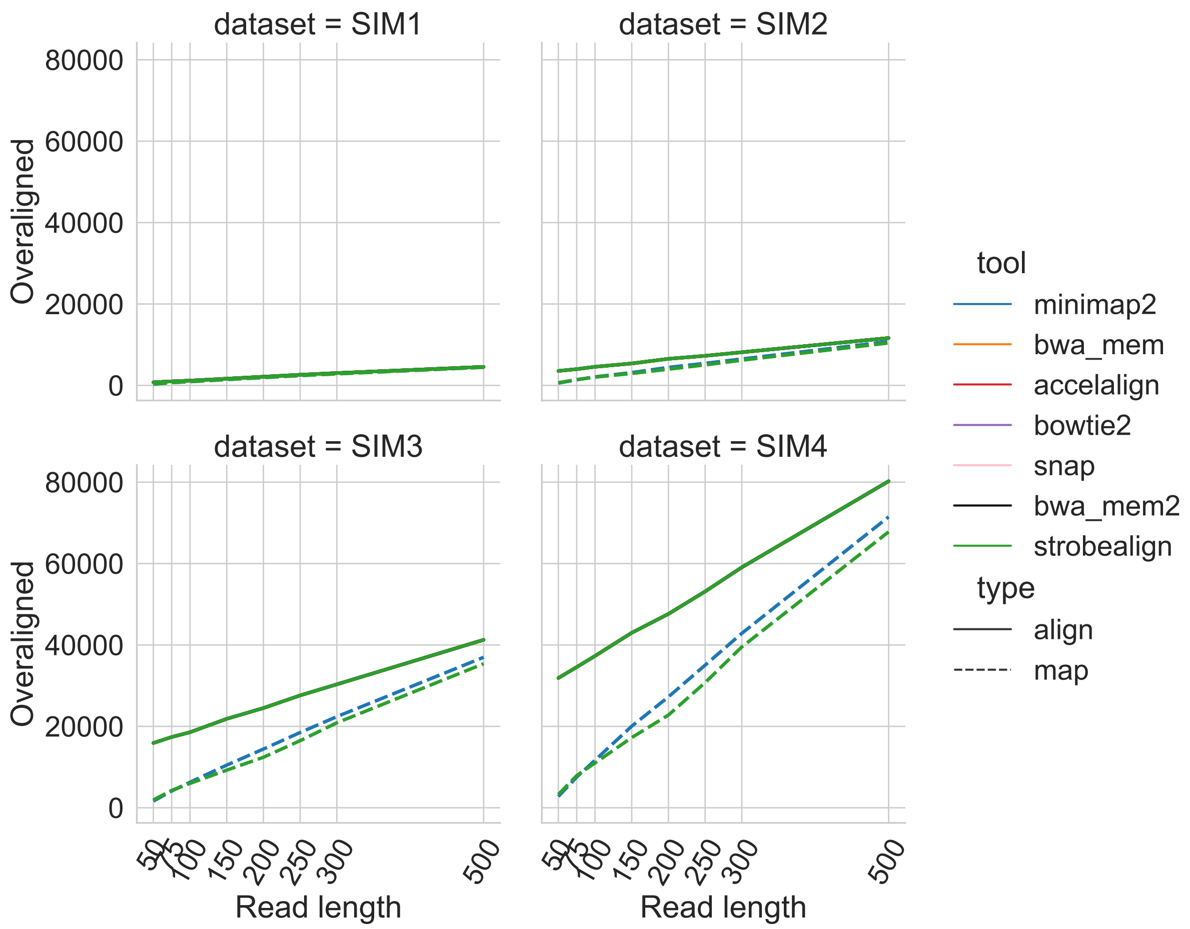


Figure S11: The number of aligned single-end reads for reads classified as unaligned by the mason simulator. All the aligners have the exact same number of over-aligned read which give plots with only one visible line.

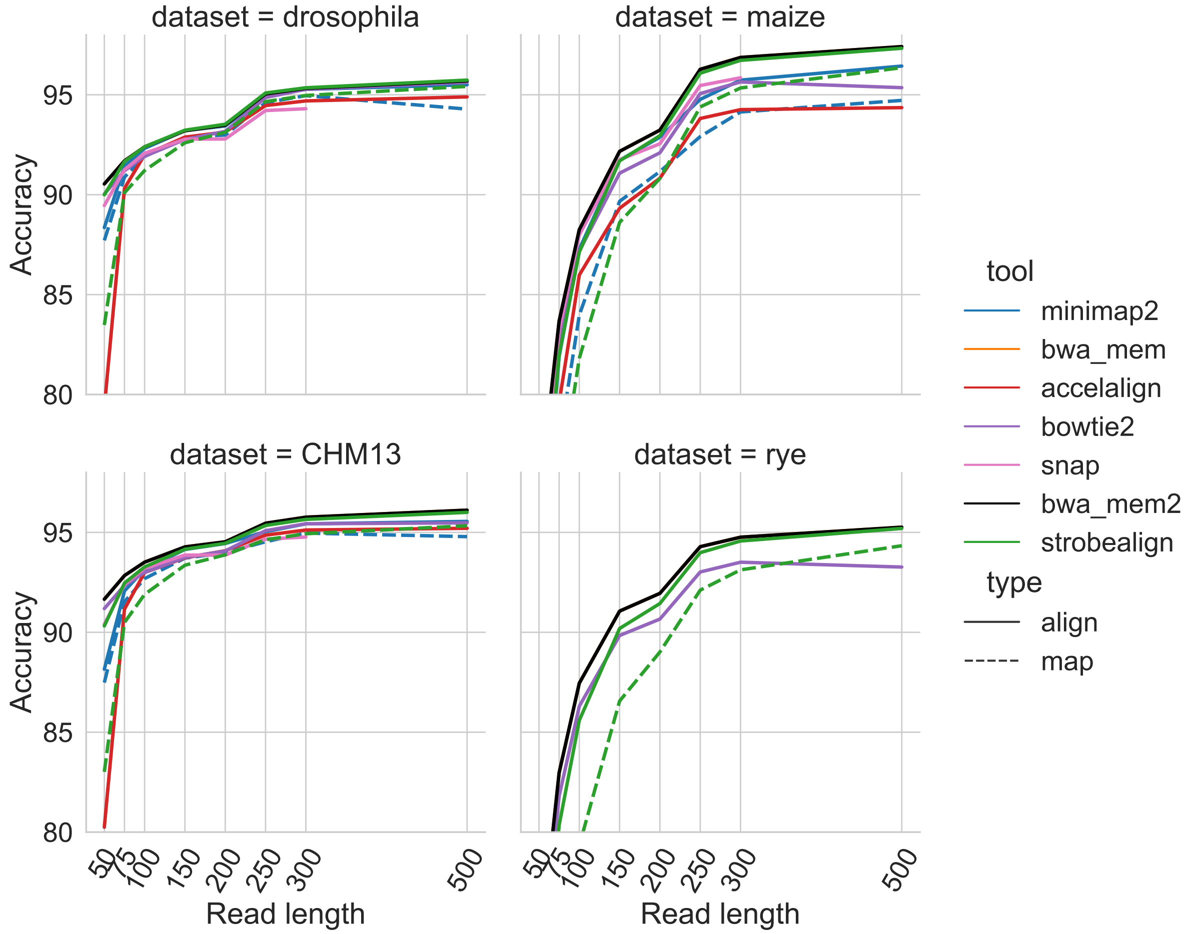


Figure S12: Alignment accuracy to the drosophila, maize, CHM13, and rye genomes. The x-axis shows simulated read length and the y-axis shows the accuracy.

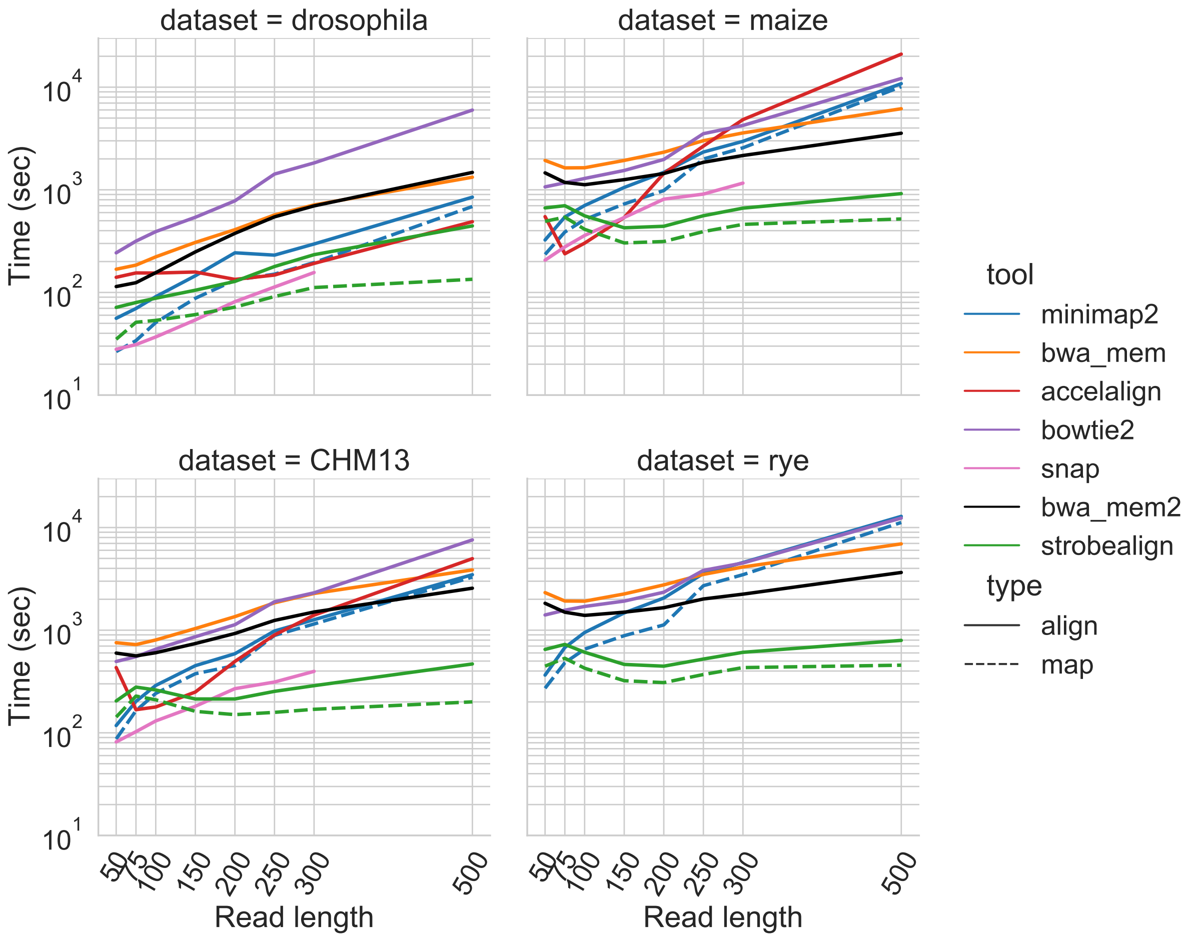


Figure S13: Alignment time for the drosophila, maize, CHM13, and rye datasets. The x-axis shows simulated read length and the y-axis shows the wall clock time from after the index has been loaded into RAM.

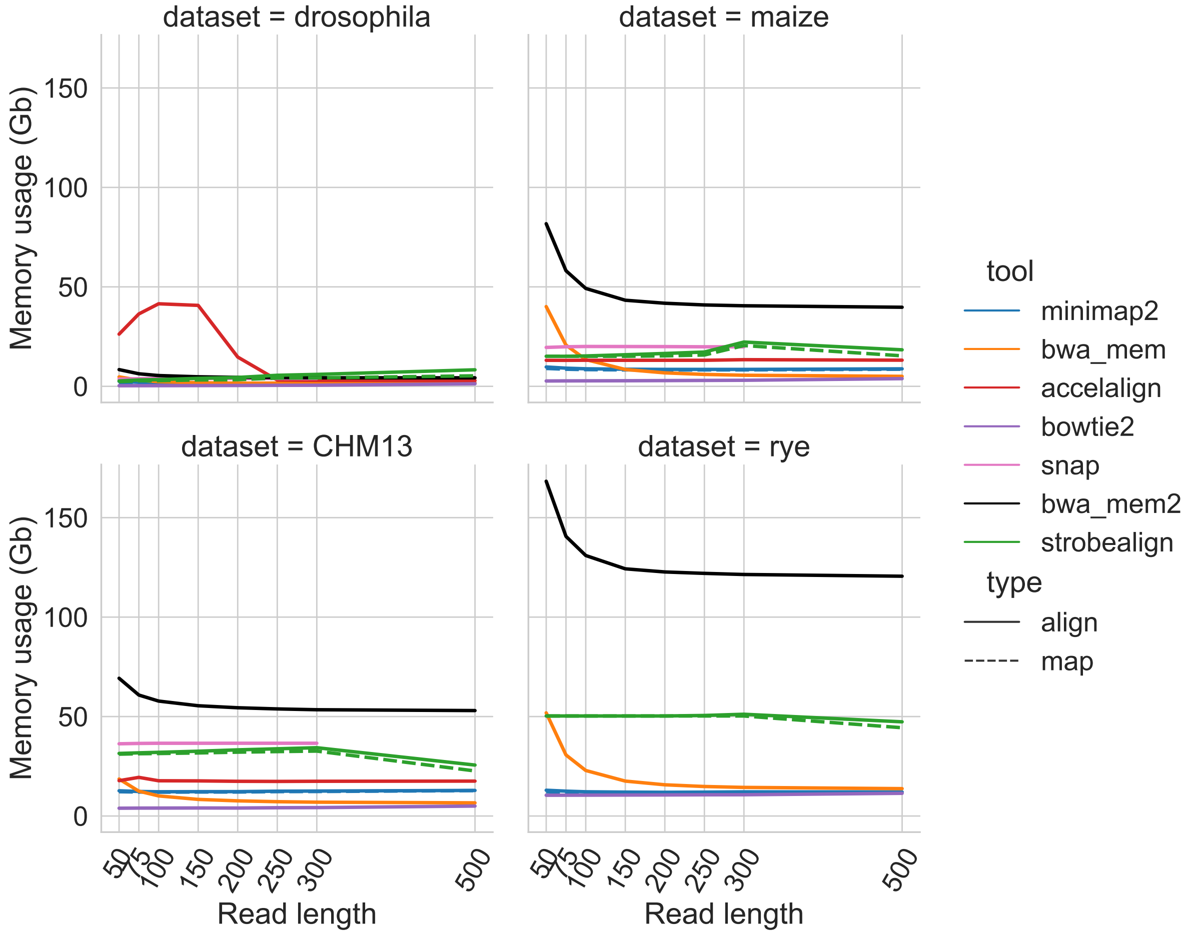


Figure S14: Peak memory usage at alignment time for the drosophila, maize, CHM13, and rye datasets.

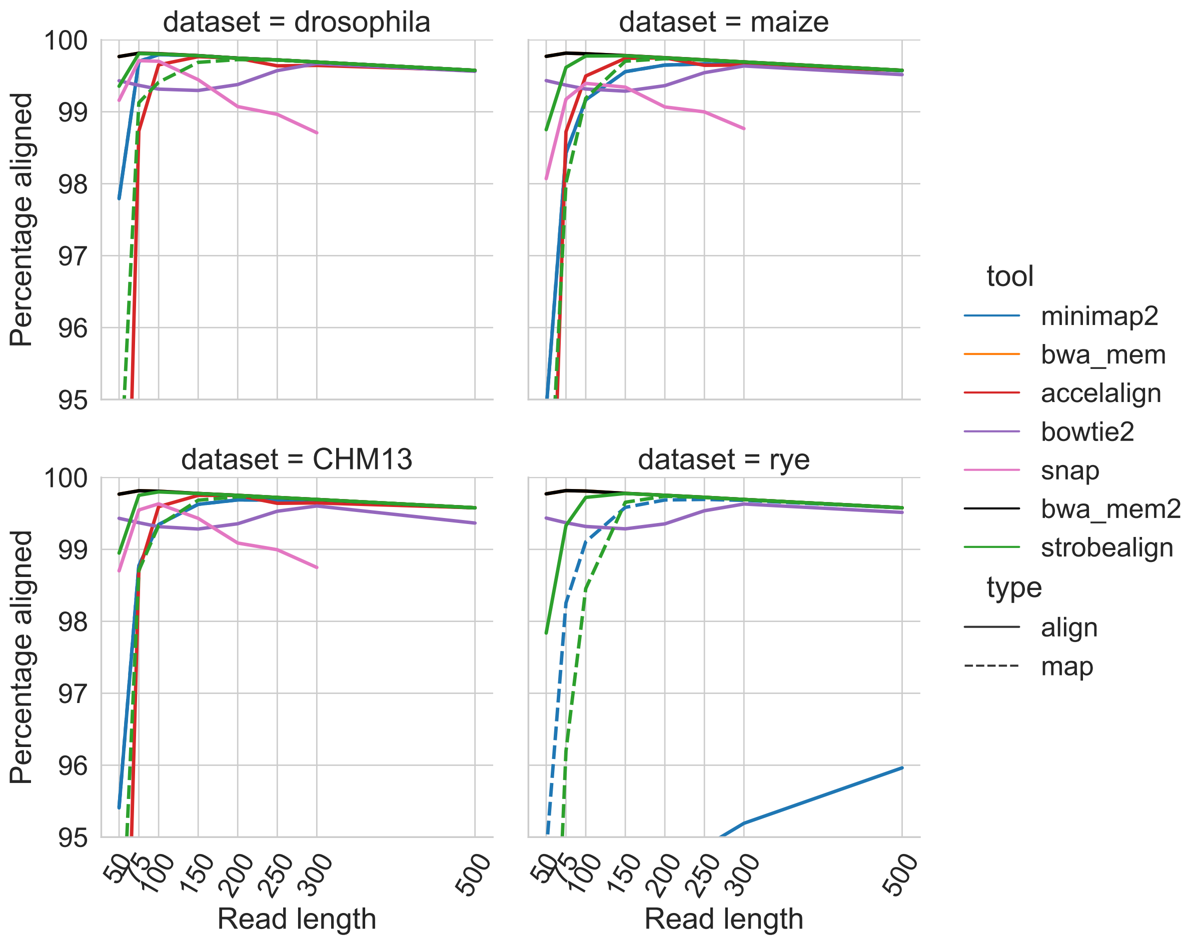


Figure S15: Percent aligned reads to the drosophila, maize, CHM13, and rye genomes. The x-axis shows simulated read length and the y-axis shows the percentage of aligned reads.

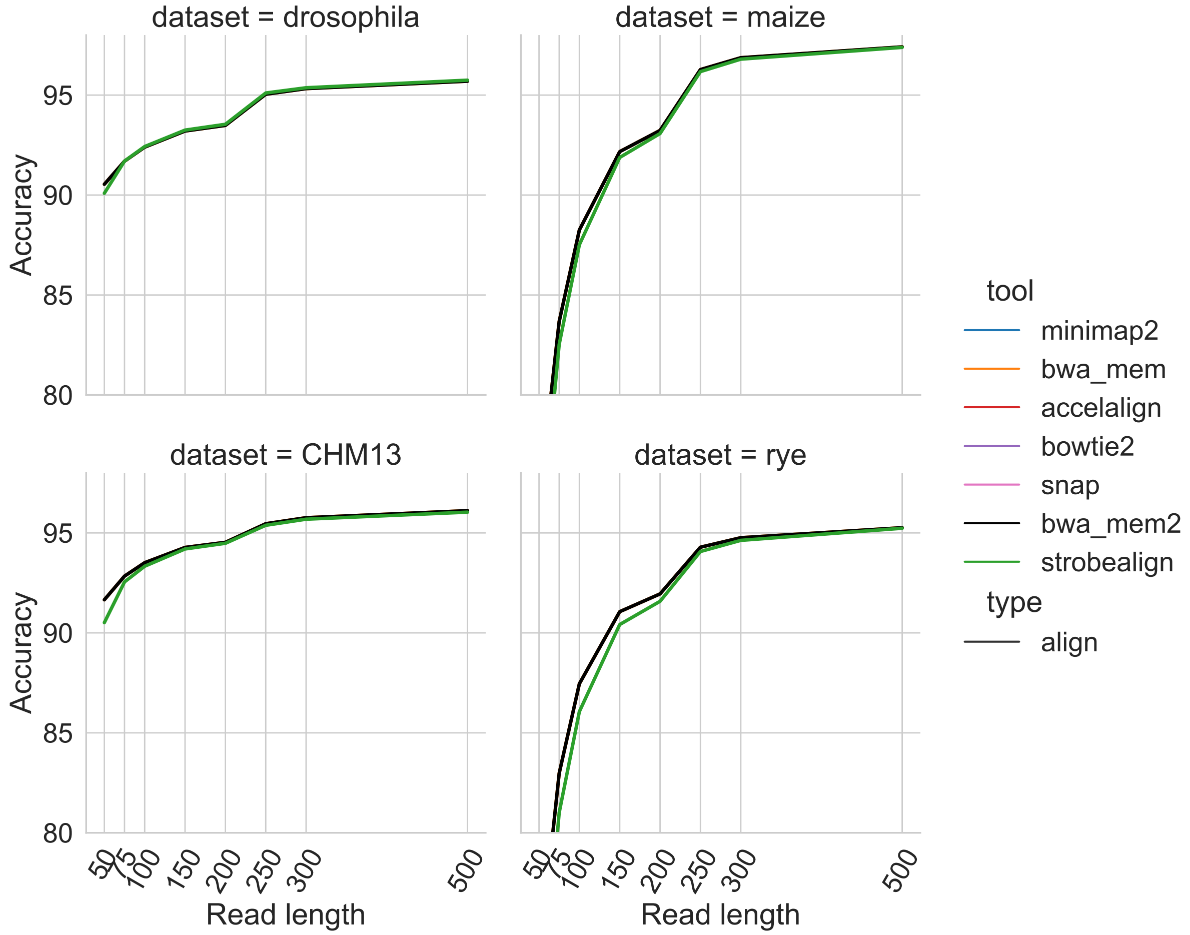


Figure S16: Alignment accuracy to the drosophila, maize, CHM13, and rye genomes when specifying -M 40 to strobealign. The x-axis shows simulated read length and the y-axis shows the accuracy.

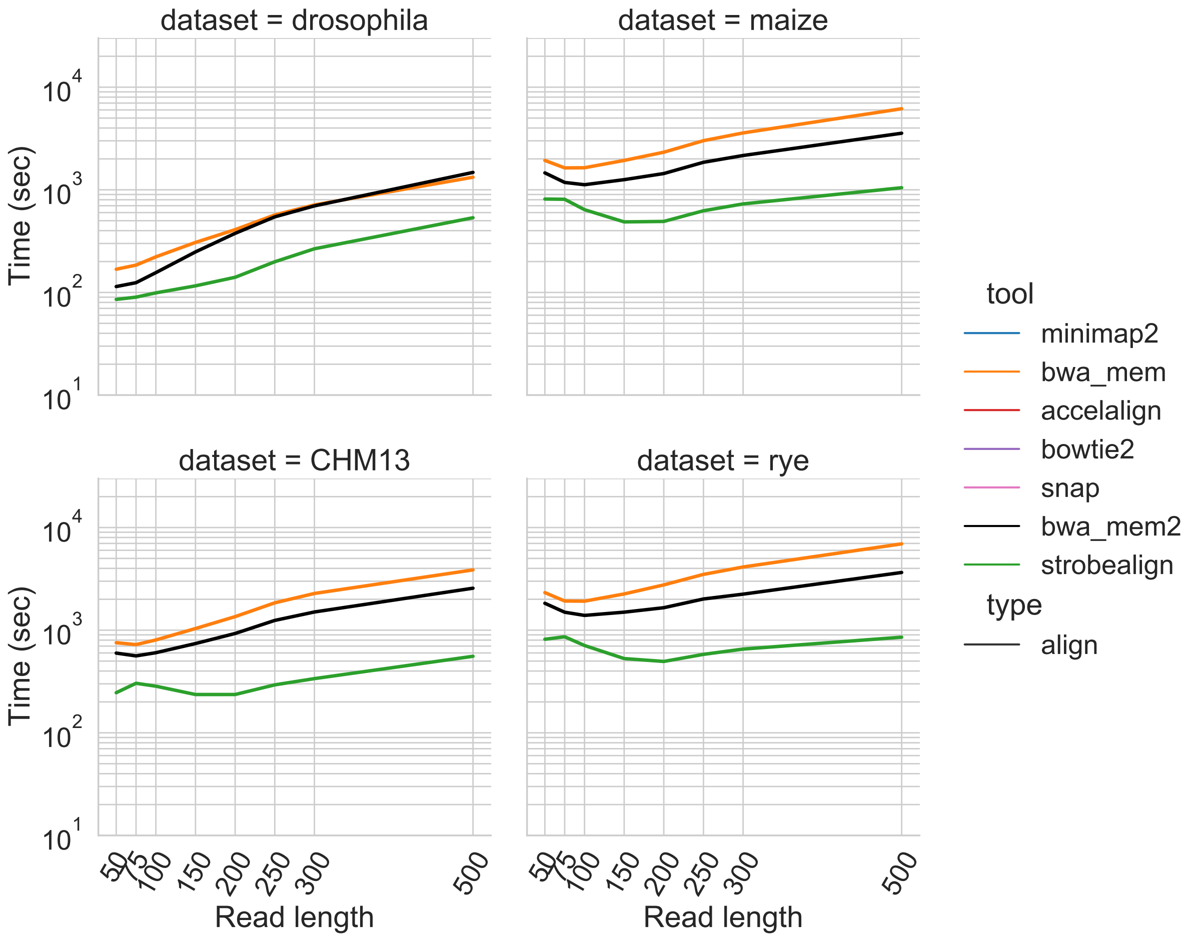


Figure S17: Alignment time for the drosophila, maize, CHM13, and rye datasets when specifying -M 40 to strobealign. The x-axis shows simulated read length and the y-axis shows the wall clock time from after the index has been loaded into RAM.

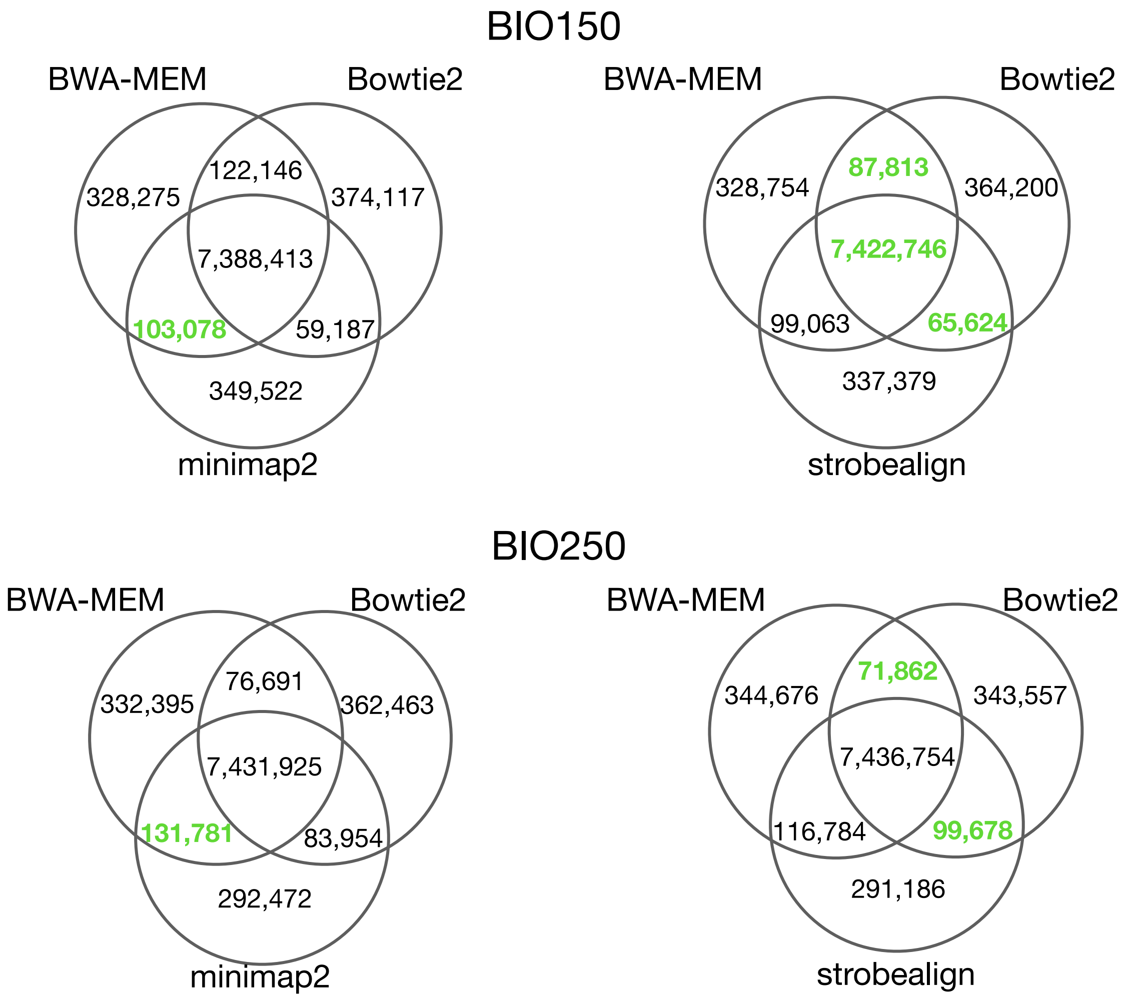


Figure S18: The number of aligned reads in the BIO150 and BIO250 datasets with genomic overlap between the two commonly used short-read aligners BWA-MEM and Bowtie2 and minimap2, and strobealign. Due to lack of ground truth, we view an overlap with either BWA-MEM or Bowtie2 as an indication of a good alignment. Under this assumption, the best result between minimap2 and strobealign is shown in green boldfaced text if it is more than 1% better than other aligner.
